# Entorhinal cortex vulnerability to human APP expression promotes hyperexcitability and tau pathology

**DOI:** 10.1101/2023.11.06.565629

**Authors:** Annie M Goettemoeller, Emmie Banks, Prateek Kumar, Viktor J Olah, Katharine E McCann, Kelly South, Christina C Ramelow, Anna Eaton, Duc M Duong, Nicholas T Seyfried, David Weinshenker, Srikant Rangaraju, Matthew JM Rowan

**Affiliations:** Department of Cell Biology, Emory University School of Medicine, Atlanta, GA, 30322; GDBBS Graduate Program, Laney Graduate School, Emory University; Department of Neurology, Yale University, New Haven, CT 06510; Department of Human Genetics, Emory University School of Medicine; Department of Neurology, Emory University School of Medicine; Department of Biomedical Engineering, Georgia Institute of Technology, Atlanta, GA 30322; Department of Biochemistry, Emory University; Center for Neurodegenerative Disease, Emory University School of Medicine

**Keywords:** APP, inhibition, PV interneuron, hyperexcitability, lateral entorhinal cortex, Alzheimer’s Disease

## Abstract

Preventative treatment for Alzheimer’s Disease is of dire importance, and yet, cellular mechanisms underlying early regional vulnerability in Alzheimer’s Disease remain unknown. In human patients with Alzheimer’s Disease, one of the earliest observed pathophysiological correlates to cognitive decline is hyperexcitability. In mouse models, early hyperexcitability has been shown in the entorhinal cortex, the first cortical region impacted by Alzheimer’s Disease. The origin of hyperexcitability in early-stage disease and why it preferentially emerges in specific regions is unclear. Using cortical-region and cell-type-specific proteomics coupled with *ex vivo* and *in vivo* electrophysiology, we uncovered differential susceptibility to human-specific amyloid precursor protein (hAPP) in a model of sporadic Alzheimer’s. Unexpectedly, our findings reveal that early entorhinal hyperexcitability may result from intrinsic vulnerability of parvalbumin (PV) interneurons, rather than the suspected layer II excitatory neurons. This vulnerability of entorhinal PV interneurons is specific to hAPP, as it could not be recapitulated with increased murine APP expression. However, partial replication of the findings could be seen after introduction of a murine APP chimera containing a humanized amyloid-beta sequence. Surprisingly, neurons in the Somatosensory Cortex showed no such vulnerability to adult-onset hAPP expression. hAPP-induced hyperexcitability in entorhinal cortex could be ameliorated by enhancing PV interneuron excitability *in vivo.* Co-expression of human Tau with hAPP decreased circuit hyperexcitability, but at the expense of increased pathological tau species. This study suggests early disease interventions targeting non-excitatory cell types may protect regions with early vulnerability to pathological symptoms of Alzheimer’s Disease and downstream cognitive decline.

## Introduction

Alzheimer’s Disease (AD) is the most prevalent neurodegenerative disease, yet current treatments are unable to prevent its initiation and progression. Although brain regions of early vulnerability have been known for over 30 years^1^, our understanding of what makes certain areas more susceptible remains unknown. The first cortical region to display pathology and degeneration in AD is the Lateral Entorhinal Cortex (LEC)^1–4^. Notably, landmark studies identified Layer II (LII) neurons cells as highly vulnerable to early neurodegeneration with up to 60% cell death in mild AD patients and up to 90% in severe cases^2^. More recently, LII LEC principal neurons were also characterized as a cell population exhibiting amyloid pathology^4^. However, the distinctive features that impart vulnerability to neurons in the LEC AD remain unclear. Uncovering region-specific cellular mechanisms could improve our understanding of the initiating factors in the AD cascade and are imperative in determining potential interventions at a time when subsequent cognitive decline and neurodegeneration might still be prevented.

Hyperexcitability is one of the earliest pathophysiological biomarkers in the human AD brain, and its emergence correlates with severity of cognitive decline in individuals^5^. Hyperexcitability is also observed in recordings from *in vivo* and *in vitro* models of AD pathology^6–12^, arising prior to amyloid plaque deposition^13^ and likely contributing to spine degeneration^14^. Interestingly, hypermetabolism^15^ and hyperexcitability^9,16^ emerged in the LEC of a sporadic AD mouse model before spreading to other regions^3^. It is unclear whether cell-intrinsic changes in principal neuron excitability or other forms of circuit dysfunction are responsible for aberrant LEC activity in early AD. Hyperexcitability may also arise due to changes in local circuit inhibition from GABAergic interneurons, with several lines of evidence demonstrating impaired inhibitory tone^6,9,15^, most notably from fast-spiking parvalbumin+ (PV) interneurons^7,10,13^. Whether the basal properties of PV interneurons in the LEC confer functional vulnerability with respect to PV cells in other regions is unknown. Thus, observing baseline cellular and regional differences coupled with adult-onset, region-specific APP or Tau expression is imperative to properly dissect inherent vulnerabilities underlying susceptibility of the LEC to early AD pathology.

## Results

### PV interneurons in an AD-vulnerable region are functionally and molecularly distinct

We first compared active and passive features of excitatory neurons in AD-vulnerable and non-vulnerable cortical regions. Excitatory neurons in LII of Lateral Entorhinal Cortex (LEC) (highly vulnerable to early AD pathology^4^) and L5 pyramidal cells (PCs) in Somatosensory Cortex (SS Ctx) of wild type (WT) mice were chosen for comparison, as each represent projection output neurons and are innervated by similar dominant inhibitory networks^17^. Despite differences in their dendritic anatomy, axonal projections, and overall local circuit operations, these two cell types showed striking overlap in their firing capacity, AP waveforms, and most other biophysical features (Fig 1a-c), with only slight biophysical differences noted (Extended Data Table 1). Because different cortical regions perform operations over non-overlapping frequency domains, we hypothesized that differences in the intrinsic excitability of inhibitory interneurons might help tune circuit activity locally. Thus, we assessed physiological phenotypes of ‘fast-spiking’ PV interneurons in each region, using an unbiased, PV-specific enhancer-AAV fluorescent targeting approach^18^. In the LEC, the E2 enhancer displayed high overlap (92.62*±*5.7%) with PV+ somas from the previously established mouse model, PV-tdTom (Extended Data Fig. 7a,b). Surprisingly, PV interneurons in the LEC maximally fired at only half the rate of SS Ctx PV interneurons (Fig. 1d,e), likely due to their far broader action potentials with respect to PV interneurons recorded from SS Ctx (Fig. 1f-h)^17^. The first action potential of each AP train was also larger in amplitude in the LEC PV interneurons (Fig. 1f-h). Furthermore, resting membrane potential and AP threshold were significantly different for PV cells when compared by region (Extended Data Table 1, bottom). Despite expressing similar passive features in LEC and SS Ctx (e.g., membrane capacitance; 70.17 ± 5.46 pF vs. 71.91 ± 9.51 pF; LEC vs SS respectively, Extended Data Table 1), their starkly divergent excitability suggests unique molecular signatures which may also underlie differential vulnerability in AD and other diseases.

**Figure 1.**
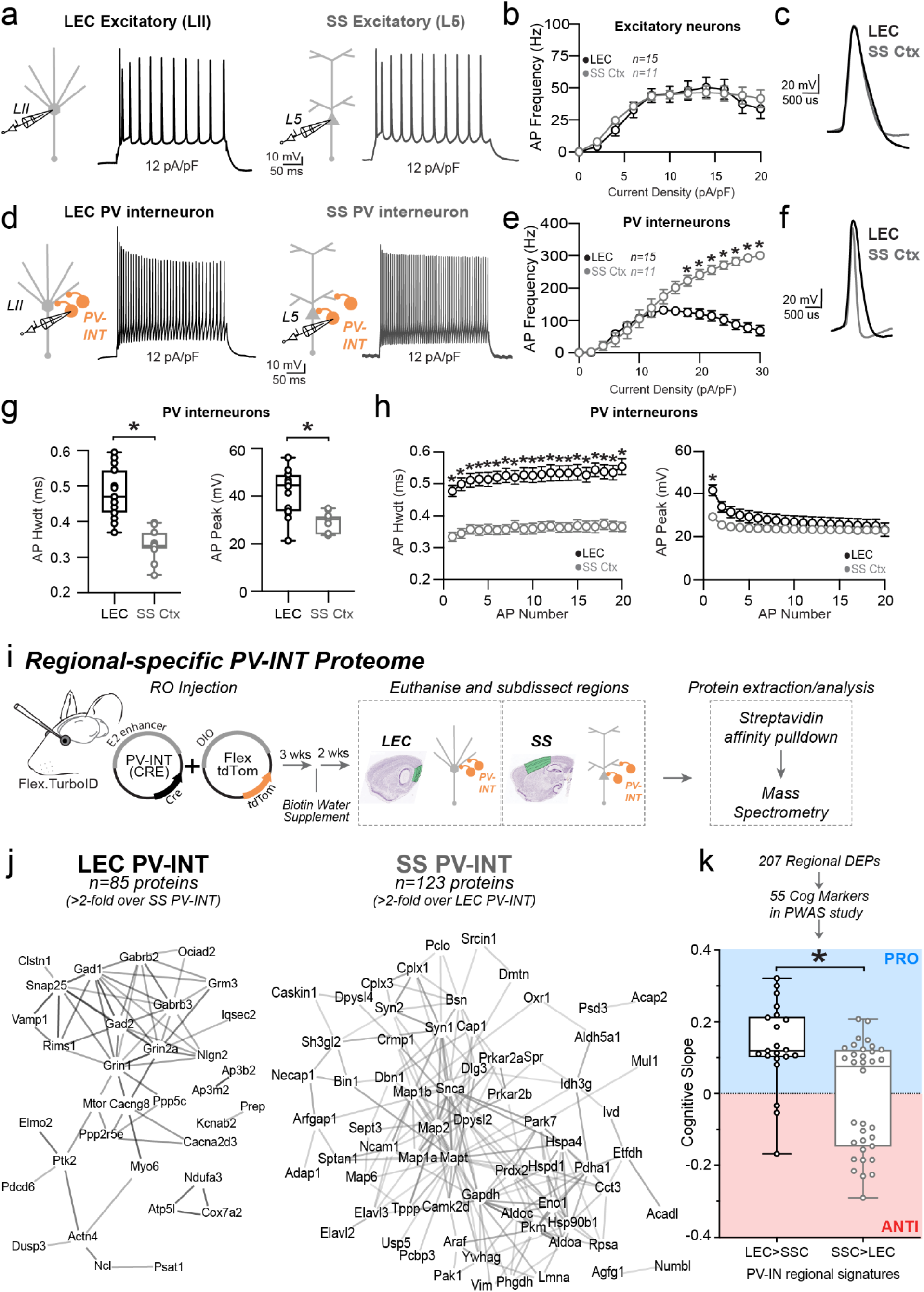
PV-INTs in an AD vulnerable region display reduced baseline firing. **a,d.** Graphical summary of AAV.E2.tdTom stereotactic injection in either the Lateral Entorhinal Cortex or Somatosensory cortex. PV-interneurons were fluorescently targeted for whole-cell current clamp recordings (d) as well as nearby excitatory cells (a). AP firing elicited by square pulse current injections of varying magnitude normalized to cellular capacitance during recording in excitatory cells (a) and PV-interneurons (d) mice from L2 LEC (left) and L5 SS Ctx (Right) at 12 pA/pF. **b.** Group data summary of AP firing frequency in WT mice. Excitatory cells between LEC and SS Ctx showed no difference in AP Frequency (Hz) Ctx (LEC: Max: 50.42±5.63 Hz, SS Ctx: Max: 46.35±5.51 Hz, p=0.46). **c.** AP waveforms of excitatory cells were compared at 12 pA/pF square pulse injections in WT mice from L2 LEC and L5 SS Ctx. Aps from the 1^st^ spike in the train are superimposed for comparison. **e.** Group data summary of AP firing frequency in WT mice. PV interneurons in L2 LEC show a strong reduction in AP max firing frequency at higher current densities when compared to PV interneurons of L5 SS Ctx (LEC: Max: 131.6±11.48 Hz, SS Ctx: Max: 301.1±27.59 Hz, p=<0.0001 for 16 pA/pF and above). **f.** AP waveforms of tdTom+ PV interneurons were compared at 12 pA/pF square pulse injections in WT mice from L2 LEC and L5 SS Ctx. Aps from the 1^st^ spike in the train are superimposed for comparison. **g.** Summary data of AP properties. L2 LEC PV interneurons display a significantly increased AP peak (LEC: 41.86±2.66 pA, SS: 28.75±1.30 pA, p=0.0002, t= 4.83, df=24, two-tailed unpaired t-test) and AP Hwdt (LEC: 0.48 ± 0.02 ms, SS: 0.33 ± 0.01 ms, p=<0.0001, t=6.10, df=25) for the first AP of the spike train. Individual data points and box plots are displayed. Significance is defined as p<0.05, unpaired t-tests. **h.** Relationship between AP peak or width, in WT mice and AP # during spike trains elicited with a 12 pA/pF current injection. **i.** Experimental approach for Regional-specific PV-interneuron Proteomes: E2 enhancer Cre AAV was retro-orbitally delivered to WT (Control) or Rosa26TurboID/wt (PV-CIBOP) mice (n=3 per genotype, including males and females) followed by 3 weeks of Cre-mediated recombination, and 2 additional weeks of biotin supplementation (drinking water). The brain was then microdissected into LEC and SS Ctx and prepared for biochemical studies. **j.** STRING analysis of PV-enriched proteins for LEC PV-INTs (left) and SS Ctx PV-INTs (right) (>2-fold enriched over other region) shows synaptic receptors, synaptic vesicle and exocytosis related proteins including GAD1/2, GABAb2/3, and complexins. **k.** Enrichment of PWAS-identified proteins associated with cognitive slope in LEC (left) or SS Ctx (right) PV-enriched proteomic signatures. Cognitive slope was estimated in ROSMAP cases. Positive slope indicates cognitive stability or resilience when proteins are present while a negative slope indicates cognitive decline when proteins are present. Proteins positively correlated with cognitive slope are referred to as pro-resilience proteins while those negative correlated with cognitive slope are anti-resilience proteins. Enrichment of previously identified ‘pro’ and ‘anti’ resilience proteins within the PV protein dataset identified by CIBOP were assessed after weighting based on strength of association between proteins and cognitive slope. (LEC: 0.13± 0.03 and SS Ctx: −0.01±0.03; p=0.001, t=3.71, df=53, two-tailed Mann Whitney test). For b, e, and h: For all summary graphs, data are expressed as mean (± SEM). Statistical significance is denoted as *=p<0.05, as determined by Two-way ANOVA with Sidak’s multiple comparison test. For all summary graphs, data are expressed as mean (± SEM). Also see Extended Data Figure 1 for related analyses and datasets.

We next sought to examine differences in the molecular signature of PV interneurons in each of these regions. Single-neuron transcriptomics is a sound method for uncovering molecular diversity between different brain cell types. Nonetheless, the functional relevance of these studies is limited by substantial discordance between mRNA and protein in neurons^19^. Thus, we opted to isolate the native-state proteomes of PV interneurons from each region using our recently developed neuron-type-specific TurboID method^20^ (Fig. 1i). This was achieved through systemic AAV injections to achieve whole-cortex expression of a PV-specific, Cre-expressing enhancer-AAV in Flex.TurboID mice^21^ followed by region-specific microdissection (Fig. 1i; Extended Data Fig. 1a). Flex.TurboID mice express an engineered biotin ligase (TurboID) in a cell-specific manner. After recombination, TurboID will efficiently label the extranuclear proteins in cell types of interest. Thus when coupled with subdissection, region- and cell-specific proteomic analyses can be performed. Over 800 proteins were biotinylated in PV interneurons in each region, of which nearly two hundred proteins showed region-specific differential abundances (unadjusted p<0.05 n=207; n=185 below the permFDR 0.05 threshold; Extended Data Datasheet 1; Fig. 1j). Generally, LEC PV interneuron proteomes showed biased enrichment in transmembrane and synaptic ion channels and transporters, while SS PV interneuron proteomes showed biased enrichment in microtubule binding, glycolysis, and fatty acid metabolism-related proteins (Extended Data Fig. 1b).

### Relationships between PV interneuron proteomic signatures with cognitive resilience in human AD

We next considered whether PV interneuron proteins differentially expressed by region (Fig. 1j) were representative of proteins associated with cognitive stability during aging. To achieve this, we used data from a protein-wide association study of cognitive resilience from human brain samples (Religious Orders Study and the Rush Memory and Aging Project; ‘ROSMAP’^22^). In this study, rate of cognitive decline (cognitive slope) was correlated with post-mortem brain protein levels quantified by mass spectrometry, identifying proteins positively associated with cognitive stability (pro-resilience proteins) and those negatively associated with cognitive stability (anti-resilience proteins) (Fig. 1k). We found that wild-type LEC PV interneurons displayed significantly more ‘pro-resilience’-associated proteins as compared to SS Ctx PV interneurons (p=0.0011; Mann-Whitney test; Fig. 1k). As nearly all the LEC PV interneuron enriched proteins were associated with cognitive stability during aging, we next explored whether expression of these enriched proteins was perturbed throughout stages of AD pathology.

We examined whether expression of these enriched proteins was perturbed throughout stages of AD pathology in humans. While several proteomics surveys of post-mortem brain tissues from AD and control brain have been performed, few studies have published data comparing the entorhinal cortex (EC) to neocortical regions, such as the frontal cortex (FC). This is true particularly regarding disease staging. In a recent study^23^, EC and FC regions from post-mortem brains of control and AD cases (BRAAK stages I-III [early] and IV-VI [late]) were analyzed by quantitative MS. This yielded 737 differentially-enriched proteins (DEPs) comparing AD to control, at either early (BRAAK I-III) or late (BRAAK IV-VI) stages, which were significant in either EC or FC regions. Among these, 93 human DEPs were observed in our PV-CIBOP proteome (Extended Data Fig. 2a, Extended Data Datasheet 2). Of these, 23 proteins showed differential levels in SS Ctx PV-INs as compared to LEC PV-INs (Extended Data Fig. 2b). Surprisingly, of the regional PV-IN proteins that were altered in human AD brain, many were pro-resilience proteins. Importantly, the LEC-enriched PV-IN proteins (including pro-resilience proteins) showed decreased levels in the EC of human AD cases (Extended Data Fig. 2b). Thus, resilience factors in PV-INs of the entorhinal cortex may be lost as AD pathology advances.

Based on observed associations between regional proteomic signatures of PV interneurons with cognitive resilience and with early changes occurring in human AD brain, we further assessed relationships between regional proteomic signatures of PV interneurons with APP and Tau protein-protein interactomes. Many APP-interacting proteins have been identified. Of 243 APP interactors identified from physical protein-protein interactors listed in the STRING database, 31 APP interactors were identified in PV-CIBOP proteomes. From these, 14 proteins were highly enriched regionally in PV interneurons. 10 proteins were highly enriched in SS Ctx PV interneurons (including Numbl, Snca, Mapt, Bin1, Hspd1, Hspa4, Hspa8, Eno1, Gapdh, Mapk3) (Extended Data Fig. 3a) and 4 were enriched in LEC PV-INs (Apoo, Grin1, Clstn1, Grin2a) (Extended Data Fig. 3a, Extended Data Datasheet 3). However, these physical associations may be altered in AD with either altered APP expression or mutations in the APP gene.

In a consensus analysis of tau protein interactors^24^, over 2000 tau protein interactors were identified across seven human post-mortem proteomics studies. Of these, 261 proteins were identified consistently as interactors (represented in at least three of the studies), comprising a high-confidence list of tau interactors. These proteins, as previously described^24^, were enriched in proteins involved in protein translation, mRNA processing and splicing, protein folding, intracellular transport, proteasome assembly, and glycolysis. In our work, 107 of these were labeled by CIBOP in PV interneurons (Extended Data Datasheet 3,4). Of these, 32 proteins had higher levels in SS Ctx PV interneurons, 8 with higher levels in LEC PV interneurons, while 67 did not show regional differences (Extended Data Fig. 3b). In contrast, non-tau interactors were more evenly distributed across SSC and LEC PV interneurons. This result suggests that tau interactors are present at higher levels in PV-INs located in the SS Ctx as compared to those in the LEC (Chi square statistic 20.7, p=0.00032). This seems to be consistent with higher levels of MAPT in SS Ctx PV interneurons as well. Tau interactors in SS Ctx PV interneurons included 14-3-3 proteins, heat shock proteins, extracellular vesicle proteins, actin/cytoskeletal proteins and RNA binding proteins. Whether the higher abundance of MAPT and tau interactors in SS Ctx PV interneurons influences the circuit’s resilience to hyperexcitability at early time points is unclear. To further explore differential responses of PV interneurons between the LEC and neocortex, we utilized a model of adult-onset induction of AD-related pathology which could be regionally and temporally controlled.

### Adult-onset human APP expression reduces PV interneuron excitability specifically in LEC

Traditional rodent models of AD express various (typically mutant) forms of hAPP (and related processing proteins), with transgene expression beginning while neuronal circuits are still maturing, and also in a brain-wide fashion. To eliminate the substantial network effects of hAPP during development^25^ and to assess inherent vulnerability of individual areas independently, we used an adult-onset, region-specific AAV approach. To explore whether differences in basal excitability and proteomic signatures of PV interneurons described early conferred region-specific vulnerability in an AD pathology context, we virally expressed wild-type hAPP in either the LEC or SS Ctx in 8-12 week old (adult) mice. Full length hAPP (hAPP 770) (NM_000484.4), an isoform showing increased expression in human AD^26,27^ was expressed using the pan-neuronal EF1a promoter (Figure 2a; AAV.Ef1a.hAPP). We assessed the impact of this hAPP isoform on PV interneurons in the LEC and SS Ctx independently after 2-3 weeks of expression.

**Figure 2.**
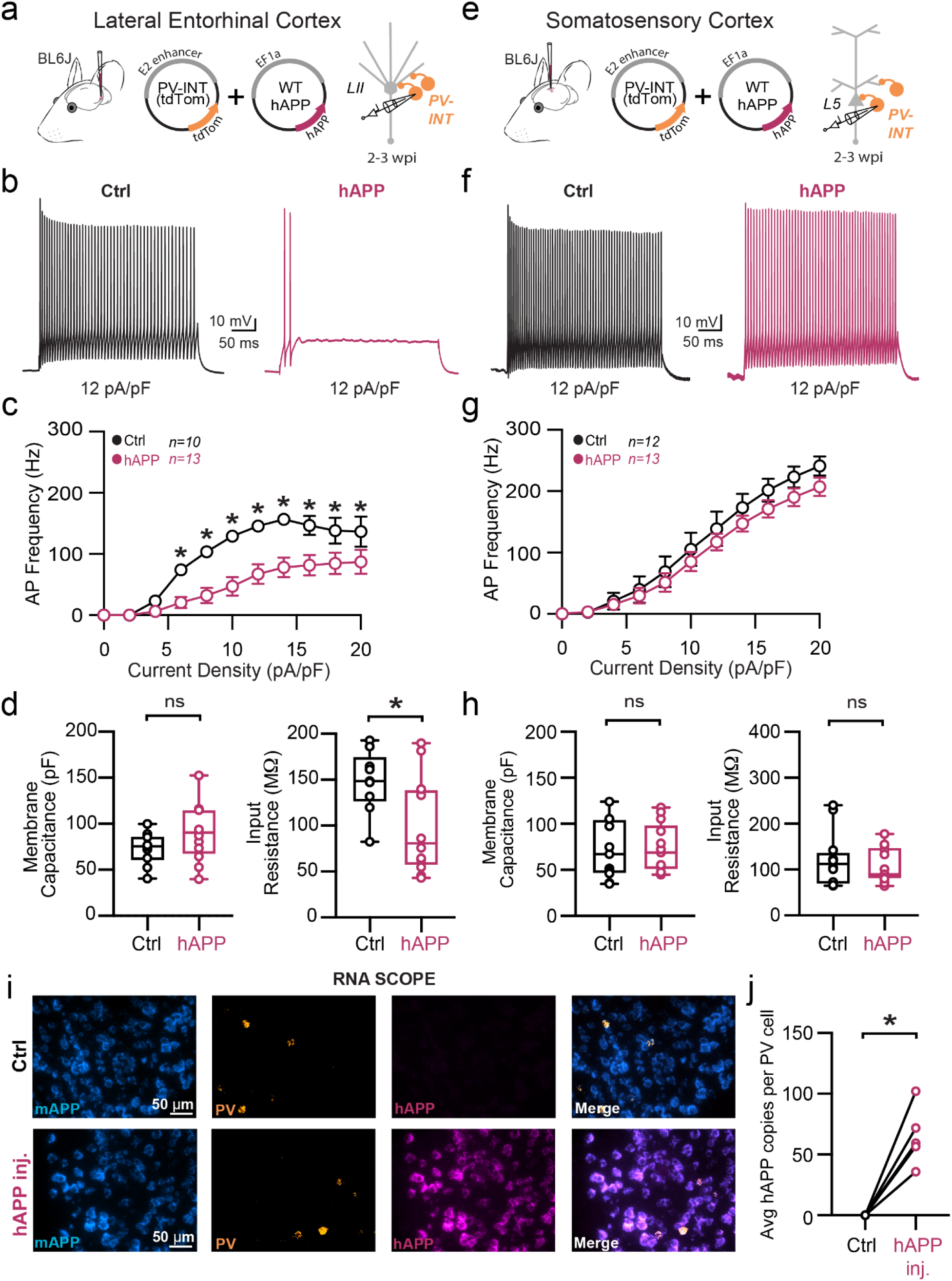
Adult-onset human APP expression reduces LEC PV interneuron excitability. **a.** Graphical summary of AAV.E2.tdTom and AAV.EF1a.hAPP (or for Ctrl, saline) stereotactic injection in the Lateral Entorhinal Cortex. PV-interneurons were fluorescently targeted (tdTom+) for whole-cell current clamp recordings. **b.** AP firing elicited by square pulse current injections of varying magnitude normalized to cellular capacitance during recording in tdTom+ PV-INT from L2 LEC at 12 pA/pF. **c.** Group data summary of AP firing frequency in L2 LEC from Ctrl (black) and hAPP injected mice (magenta). LEC PV interneurons from hAPP injected mice show a significant reduction in AP Frequency (Hz) when compared to Ctrl(Ctrl: Max: 156.6 ±13.52 Hz, hAPP: Max: 91.84± 8.74 Hz). **d.** Summary data of AP properties. L2 LEC PV interneurons after hAPP injection display a significantly decreased input resistance (Ctrl: 145.7 ± 11.61 MΩ, hAPP: 88.78 ± 15.11 MΩ, p=0.01, t=2.73, df=21) and an insignificant increase in membrane capacitance (Ctrl: 68.83 ± 5.34 pF, hAPP: 90.21 ± 9.77 pF, p=0.07, t=1.92, df=21). **e.** Graphical summary of AAV.E2.tdTom and AAV.EF1a.hAPP (or for Ctrl, saline) stereotactic injection in the Somatosensory Cortex. PV-interneurons were fluorescently targeted (tdTom+) for whole-cell current clamp recordings. **f.** AP firing elicited by square pulse current injections of varying magnitude normalized to cellular capacitance during recording in tdTom+ PV-INT from L5 SS Ctx at 12 pA/pF. **g.** Group data summary of AP firing frequency in L5 SS Ctx from Ctrl (black) and hAPP injected mice (magenta). SS Ctx PV interneurons from hAPP injected mice show no significant change in AP Frequency (Hz) when compared to Ctrl (Ctrl: Max: 301.1 ±27.59 Hz, hAPP: Max: 257.2± 24.06 Hz). **h.** Summary data of AP properties. SS Ctx interneurons after hAPP injection display an unchanged Membrane Capacitance (Ctrl: 71.91 ± 9.514, hAPP: 73.14 ± 7.327, p=0.9180, t=0.1041, df=23) and input resistance (Ctrl: 121.2 ± 17.14, hAPP: 109.1 ± 10.56, p=0.5475, t=0.6106, df=23). **i.** RNAscope representative images at 40x magnification for Ctrl injected (top) and hAPP injected mice (bottom): mAPP mRNA (cyan), Parvalbumin mRNA (gold), human APP mRNA (magenta), and a final merged image. **j.** RNAscope quantification for hAPP copies per PV+ cell comparing control to hAPP injected. hAPP injected show a significant increase in hAPP copies per PV+ cell (p=0.0039, t=5.987, df=4; two-tailed paired t-test). For all summary graphs, data are expressed as mean (± SEM). For c, g, and i: Statistical significance is denoted as *=p<0.05, as determined by Two-way ANOVA with Sidak’s multiple comparison test. For d, h: Individual data points and box plots are displayed. Statistical significance is denoted as *=p<0.05, as determined by two-tailed unpaired t-test.

In LEC PV interneurons, we observed a dramatic reduction in PV interneuron firing (Fig. 2b,c) potentially related to a reduction in input resistance (Fig. 2d), as no other relevant factors (e.g., AP waveform, RMP, AP threshold, Membrane capacitance) (Fig. 2d; Extended Data Fig. 4) were affected. Using unsupervised clustering of LEC ‘fast-spiking’ interneuron biophysical features, control- and hAPP-expressing PV interneurons clustered separately (Extended Data Fig. 12b). By contrast, PV interneurons in the SS Ctx displayed no change in firing rate (Fig. 2g) despite an increase in the AP AHP (Extended Data Fig. 5d). All other active and passive features were unchanged in SS Ctx (Fig. 2h; Extended Data Fig. 5b-e). The presence of hAPP mRNA and protein was confirmed in PV neurons 2-3 weeks after viral injection (Fig. 2i-j; Extended Data Fig. 6) using RNAscope and PV-specific flow cytometry respectively. Together, the intrinsic excitability of PV interneurons was significantly reduced in the LEC, but not SS Ctx, following hAPP expression.

### Adult-onset murine APP expression does not affect PV interneuron physiology

Several studies of different mouse models of APP-related pathology report altered intrinsic excitability in GABAergic intereneurons^7–10,13^. Whether this is simply a result of hAPP overexpression^28^ during development or effects of its downstream cleavage products remain controversial. To address this, we next injected a virus containing full-length murine APP (mAPP) (NM_001198823.1) (Fig. 3a; AAV.Ef1a.mAPP) into the LEC. Despite a significant increase of mAPP expression over endogenous background levels (Extended Data Fig. 8d,e), changes in PV interneuron firing and input resistance seen following hAPP expression (Fig. 2) were absent following 2-3 weeks of viral mAPP expression (Fig. 3b,c; Extended Data Fig. 8b,c). Importantly, RNAscope studies confirmed that the magnitude of AAV-induced mAPP expression was similar to that of hAPP in earlier experiments (Figure 3d), indicating that the differential physiological effects were not due to variability in APP expression levels. Furthermore, as we saw no alterations in the somatosensory cortex PV-interneurons after hAPP expression, we conclude this differing response is not due to a specific inflammatory effect of the human protein alone. Thus, hAPP-induced dysfunction of LEC PV interneurons cannot be explained by over-expression of APP alone.

**Figure 3.**
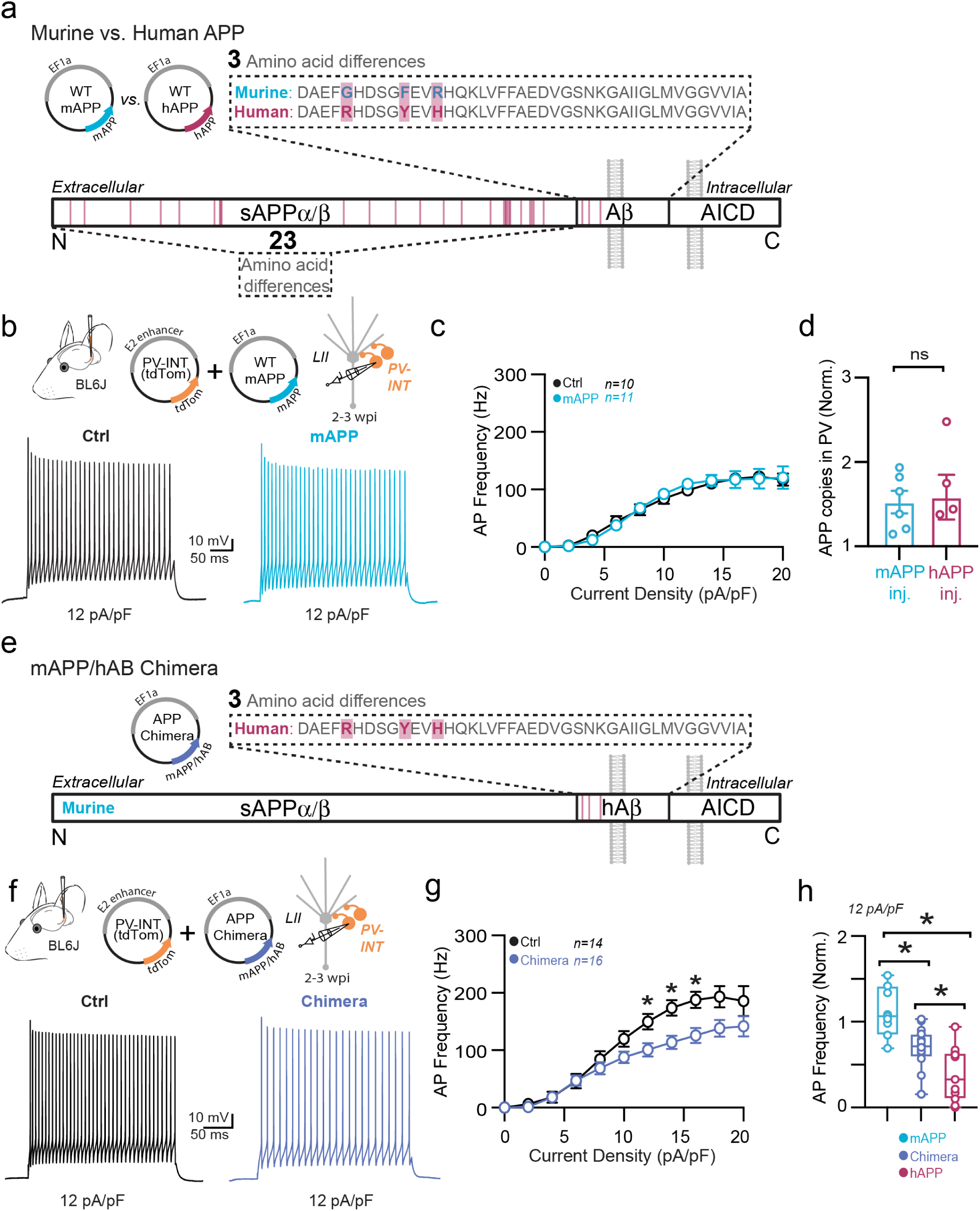
Murine APP does not affect PV interneuron physiology, but mAPP/hAB chimera replicates partial findings of hAPP-induced deficits. **a.** Pictorial representation of differing amino acids between murine APP and human APP proteins; 26 different amino acids in total, 3 of which are in the amyloid-beta segment of the protein. **b.** Graphical summary of AAV.E2.tdTom and AAV.EF1a.mAPP (or for Ctrl, saline) stereotactic injection in the Lateral Entorhinal Cortex. PV interneurons were fluorescently targeted for whole-cell current clamp recordings. AP firing elicited by square pulse current injections of varying magnitude normalized to cellular capacitance during recording in PV interneurons from Ctrl (left) and mAPP injected (right) L2 LEC at 12 pA/pF. **c.** Group data summary of AP firing frequency in Ctrl and mAPP injected mice. PV interneurons between Ctrl and mAPP injected showed no difference in AP Frequency (Hz) (Ctrl: Max: 122.3 ±11.11 Hz, mAPP: Max: 120.6 ± 11.50 Hz, p=0.95). Statistical significance is denoted as *=p<0.05, as determined by Two-way ANOVA with Sidak’s multiple comparison test. **d.** RNAscope quantification for APP copies per PV+ cell with APP injected (mAPP or hAPP) each normalized to their contralateral hemisphere average endogenous murine APP copy per PV+ cell. mAPP injected and hAPP injected mice show similar increases in increased APP expression. copies per PV+ cell (p=0.84, t=0.21, df=9; two-tailed unpaired t-test). **e.** Pictorial representation of the resultant Chimera protein; murine APP with a humanized amyloid-beta segment. **f.** Graphical summary of AAV.E2.tdTom and AAV.EF1a.mAPP/hAB Chimera (or for Ctrl, saline) stereotactic injection in the Lateral Entorhinal Cortex. PV interneurons were fluorescently targeted for whole-cell current clamp recordings. AP firing elicited by square pulse current injections of varying magnitude normalized to cellular capacitance during recording in PV interneurons from Ctrl (left) and Chimera injected (right) L2 LEC at 12 pA/pF. **g.** Group data summary of AP firing frequency in Ctrl and Chimera injected mice. PV interneurons between Ctrl and mAPP injected showed no difference in AP Frequency (Hz) (Ctrl: Max: 193.6 ±19.47 Hz, mAPP: Max: 145.4 ± 14.05 Hz, p<0.0001; for 12 pA p=0.0378, for 14 pA p=0.0368, for 16 pA p=0.0426). Statistical significance is denoted as *=p<0.05, as determined by Two-way ANOVA with Sidak’s multiple comparison test. **h.** Comparison of PV interneuron firing frequencies expressing mAPP, mAPP/hAB Chimera, or hAPP normalized to their dataset controls at 12 pA/pF. Statistical significance is denoted as *=p<0.05, as determined by Ordinary one-way ANOVA with Tukey’s multiple comparisons test. (mAPP vs. Chimera: p=0.0011, mAPP vs. hAPP: <0.0001, Chimera vs. hAPP: p=0.0335; df=35).

### Murine APP with a humanized amyloid-beta sequence moderately impairs PV interneurons

We next investigated whether the effect of hAPP on PV interneuron function could be explained by the differences in the amyloid-beta sequence of the human and mouse genes. To address this, we cloned the full-length murine APP (mAPP) (NM_001198823.1) but humanized the three differing amino acids in the amyloid-beta sequence, G676R, F681Y, and R684H (Fig. 3e; AAV.Ef1a.mAPP/hAβ Chimera). After AAV-directed expression of this APP Chimera in the LEC for 2-3 weeks, we observed a reduction in PV interneuron firing (Fig. 3f,g). However, the mAPP/hAβ effect was less robust when compared to the fully human version (Fig. 3h; Extended Data Fig. 9d) and appeared to manifest in a mechanistically distinct fashion relative to hAPP (Extended Data Fig. 9). Thus, the contribution of the human amyloid-beta sequence leads to PV-interneuron hypofunction, but was not sufficient to induce the more drastic reduction seen with full-length human hAPP. Finally, to further evaluate a role Aβ in this process, we tested the effect of exogenous Aβ (Aβ_1-40,1-42_; 0.25μM; 3 hours) incubation on PV interneurons in the LEC. Similar to hAPP, Aβ incubation alone also induced a robust reduction in AP firing (Extended Data Fig. 9f).

### Adult-onset human APP expression does not affect excitatory cell intrinsic properties

Because recent studies using different mouse models of APP/Aβ pathology report altered intrinsic excitability of excitatory neurons^25,29^, we also assessed the effects of 2-3 weeks of hAPP expression on principal excitatory cells in the LEC and SS Ctx (Fig. 4a,e). Consistent with unaltered PV firing in SS Ctx, no change in intrinsic firing frequency or passive properties were noted in pyramidal cells in the SS Ctx (Fig. 4f-h; Extended Data Fig. 11). Surprisingly, we also observed no impact of hAPP on intrinsic AP firing of LII LEC excitatory neurons (Fig. 4b,c). Further, membrane capacitance was unperturbed (Fig. 4d) suggesting no major alterations to LII cellular morphology. A modest but significant increase in dV/dt max was noted in LEC LII principal cells (Extended Data Fig. 10d), potentially via an hAPP-dependent modulation of Na_v_ channels in these cells. All other active and passive properties were unaltered (Fig. 4d; Extended Data Fig. 8). Importantly, RNAscope experiments confirmed increased hAPP expression in CaMKIIa+ cells (Fig. 4i-j), indicating that our AAV also targeted excitatory neurons as expected. Using principal component analysis (PCA) of several excitatory cell biophysical features from LEC recordings, clusters could be separated based on input resistance, membrane time constant, and resting membrane potential. These clusters likely arise due to sampling of both LII fan cells and LII pyramidal cells^30^, suggesting our population of principal cells likely included both cell types (Extended Data Fig. 12a). When assessed, these excitatory populations showed no differential clustering when expressing hAPP (Extended Data Fig. 12a). Together, these results indicate that principal neurons are more resistant to changes in their intrinsic excitability following adult-onset hAPP expression compared to PV interneurons.

**Figure 4.**
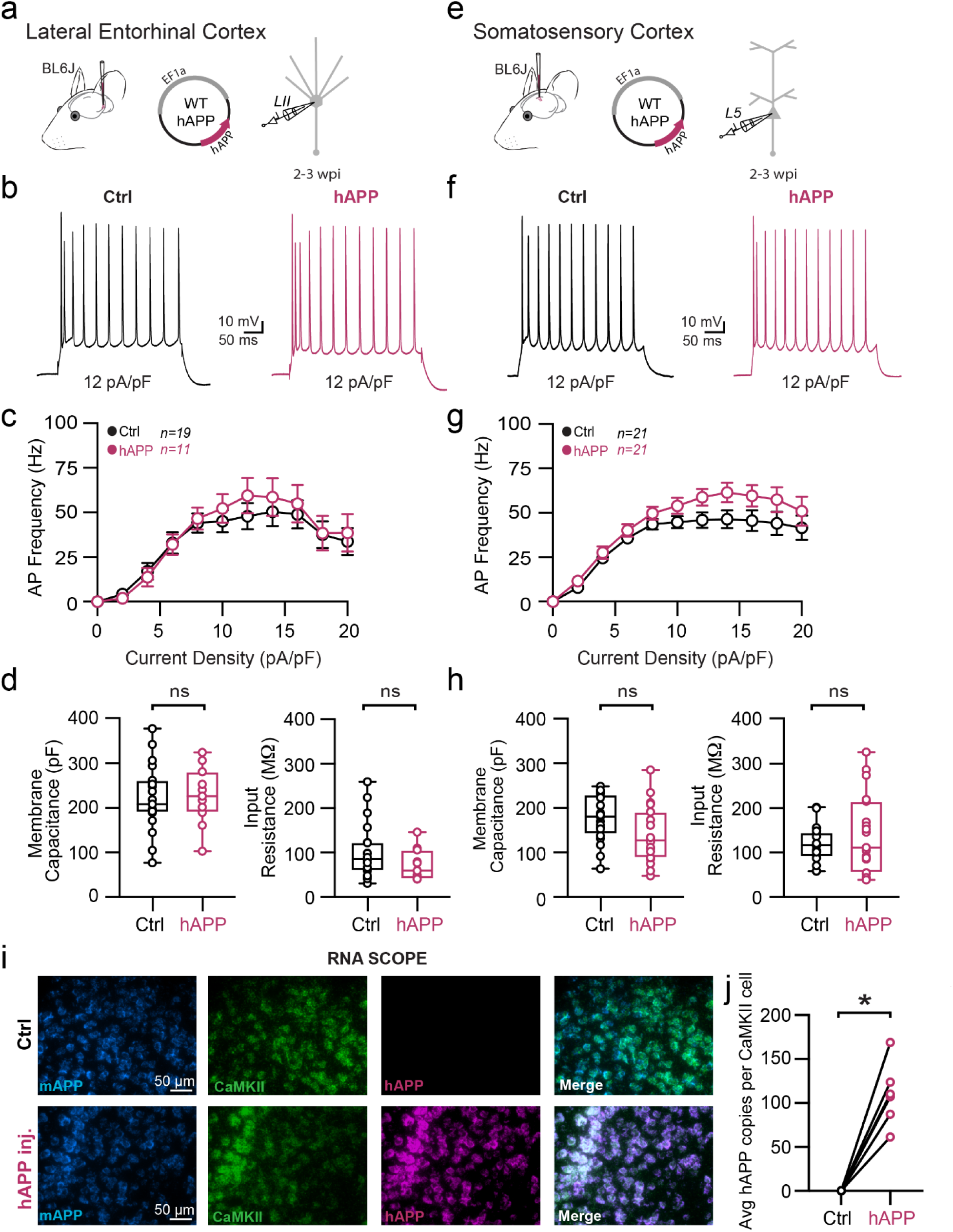
Adult-onset human APP expression does not alter excitatory neuron physiology. **a.** Graphical summary of AAV.EF1a.hAPP (or for Ctrl, saline) stereotactic injection in the Lateral Entorhinal Cortex. Excitatory cells were targeted for whole-cell current clamp recordings. **b.** AP firing elicited by square pulse current injections of varying magnitude normalized to cellular capacitance during recording in Ctrl and hAPP injected L2 LEC excitatory cells from at 12 pA/pF. **c.** Group data summary of AP firing frequency in L2 LEC from Ctrl (black) and hAPP injected mice (magenta). Excitatory neurons in L2 LEC from hAPP injected mice show no alteration in AP Frequency (Hz) when compared to Ctrl (Ctrl: Max: 50.42 ± 5.63 Hz, hAPP: Max: 59.43 ± 6.56 Hz, p=0.99, df=28). **d.** Summary data of AP properties. L2 LEC excitatory cells after hAPP injection display an unchanged Membrane Capacitance (p=0.83, t=0.27) as well as an unchanged input resistance (p=0.15, t=1.50, df=28). **e.** Graphical summary of AAV.EF1a.hAPP (or for Ctrl, saline) stereotactic injection in the Somatosensory Cortex. Excitatory neurons in L5 were targeted for whole-cell current clamp recordings. **f.** AP firing elicited by square pulse current injections of varying magnitude normalized to cellular capacitance during recording in excitatory cells from L5 SS Ctx at 12 pA/pF. **g.** Group data summary of AP firing frequency in L5 SS Ctx from Ctrl (black) and hAPP injected mice (magenta). SS Ctx excitatory neurons from hAPP injected mice show no significant change in AP Frequency (Hz) when compared to Ctrl (Ctrl: Max: 46.35 ± 5.38 Hz, hAPP: Max: 61.43 ± 6.78 Hz, p>0.05, df=40). **h.** Summary data of AP properties. SS Ctx interneurons after hAPP injection display an unchanged Membrane Capacitance and input resistance (Ctrl: 176.9 ± 11.58, hAPP: 140.5 ± 14.31, p=0.06, t=1.98, df=40, two-tailed unpaired t-test). **i.** RNAscope representative images at 40x magnification for Ctrl injected (top) and hAPP injected mice (bottom(: mAPP mRNA (cyan), CaMKIIa mRNA (green), human APP mRNA (magenta), and a final merged image. **j.** RNAscope quantification for hAPP copies per CaMKIIa+ cell comparing control to hAPP injected. hAPP injected show a significant increase in hAPP copies per CaMKIIa+ cell (p=0.0007, t=7.42, df=5; two-tailed paired t-test). For all summary graphs, data are expressed as mean (± SEM). For c, g: Statistical significance is denoted as *=p<0.05, as determined by Two-way ANOVA with Sidak’s multiple comparison test. For d, h: Individual data points and box plots are displayed. Statistical significance is denoted as *=p<0.05, as determined by two-tailed unpaired t-test.

### hAPP expression induces basal circuit hyperexcitability in the LEC but not SS Cortex

Although we observed no alterations in the intrinsic excitability of excitatory cells in either region following hAPP expression, we wanted to assess whether the changes in PV interneuron biophysics in LEC had an impact on local circuit activity. To examine this at population level, we acquired spontaneous post-synaptic currents from principal cells in either region (Fig. 5a,d). In the LEC, spontaneous inhibitory event (sIPSC) frequency was significantly decreased (increase in the mean inter-event interval [IEI]) after 2-3 weeks of hAPP expression (Fig. 5b,c). Furthermore, we analyzed the LEC sIPSCs for differences in the frequency in small and large amplitude events (cutoff 40 pA derived from a previously published method^31^), to determine if the increase in sIPSC IEI was related to distal inhibition (small amplitude) or proximal, peri-somatic inhibition (large amplitude). We observed that while the frequency of small amplitude events was unchanged (p=0.52, two-tailed unpaired t-test, t=0.65, df=18; Ctrl: 1.66±0.36 Hz, hAPP: 1.33±0.35 Hz), the frequency of large amplitude events was significantly decreased in the LEC (p=0.02, two-tailed unpaired t-test, t=2.51, df=18; Ctrl: 4.66±1.1 Hz; hAPP: 1.74±0.49 Hz), indicating a selective effect on peri-somatically innervating inhibitory neurons. In layer II of the entorhinal cortex, Reelin+ excitatory cells receive peri-somatic inhibition primarily from PV interneurons, rather than CCK basket cells^17,32^. Thus these results are consistent with a reduction in intrinsic PV excitability. In an apparent response to this reduced inhibitory tone, spontaneous excitatory event (sEPSC) frequency increased in the LEC following hAPP expression (Fig. 5b,c). In contrast to the LEC, recordings from SS Ctx (Fig. 5d,e) revealed no change in sIPSC or sEPSC frequency following hAPP expression (Fig. 5f), in agreement with the lack of changes in intrinsic excitability in the SS Ctx shown earlier. Spontaneous and miniature (excitatory or inhibitory) synaptic amplitudes in the LEC and SS Ctx were unchanged in either region (Extended Data Fig. 13), indicating that postsynaptic receptor alterations did not arise in excitatory neurons following short-term adult-onset hAPP expression. mIPSC and mEPSC frequencies were also unaltered, suggesting no change in the number of inhibitory or excitatory synapses at this point (Extended Data Fig. 13b). Together, these results indicate that basal circuit activity in the LEC, but not SS Ctx, becomes hyperexcitable following short-term hAPP expression, likely resulting from a region-specific PV interneuron vulnerability.

**Figure 5.**
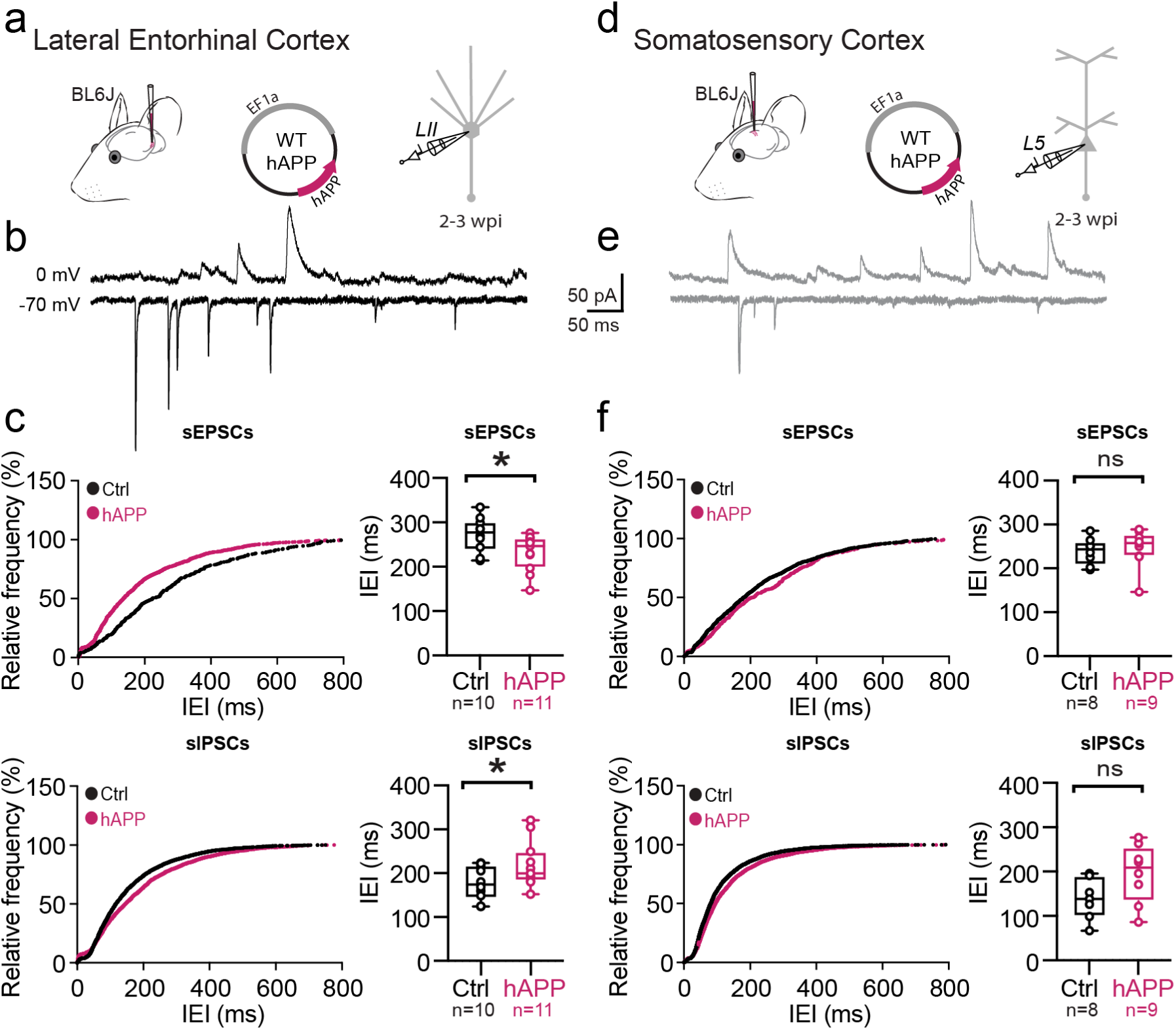
Human APP expression induces hyperexcitability in the LEC but not SS Ctx. **a.** Graphical summary of AAV.EF1a.hAPP (or for Ctrl, saline) stereotactic injection in the Lateral Entorhinal Cortex. Excitatory cells were targeted for whole-cell voltage-clamp recordings. **b.** Spontaneous events obtained by holding cell voltage at 0 mV (inhibitory post-synaptic currents, IPSCs [top]) and −70 mV (excitatory post-synaptic currents, EPSCs [bottom]), interleaved. **c. Top:** Cumulative distribution curve for spontaneous EPSCs in the LEC showing the relationship of relative frequency of events to the inter-event interval (IEI) (left). Quantified averages of IEIs are displayed for each cell as individual data points and compared between Ctrl (black) and hAPP injected (magenta) conditions (right). L2 LEC sEPSCs show a significant reduction in the IEIs (231.7 ± 12.25 ms, 272.7 ± 12.24 ms, hAPP and Ctrl respectively, p=0.029, t=2.361, df=19, two-tailed unpaired t-test). See Extended Data Fig. 13 for mEPSC data. **Bottom:** Cumulative distribution curve for spontaneous IPSCs in the LEC showing the relationship of relative frequency of events to the inter-event interval (left). Quantified averages of IEIs are displayed for each cell as individual data points and compared between Ctrl (black) and hAPP injected (magenta) conditions (right). L2 LEC sIPSCs show a significant increase in the IEIs (219.9 ± 15.84 ms, 177.3 ± 12.02 ms, hAPP and Ctrl respectively, p=0.047, t=2.097, df=19, two-tailed unpaired t-test). See Extended Data Fig. 13 for mIPSC data. **d.** Graphical summary of AAV.EF1a.hAPP (or for Ctrl, saline) stereotactic injection in the Somatosensory Cortex. Excitatory cells were targeted for whole-cell voltage-clamp recordings. **e.** Spontaneous events obtained by holding cell voltage at 0 mV (IPSCs [top]) and −70 mV (EPSCs [bottom]), interleaved. **f. Top:** Cumulative distribution curve for spontaneous EPSCs in the SS Ctx showing the relationship of relative frequency of events to the IEIs (left). Quantified averages of IEI are displayed for each cell as individual data points and compared between Ctrl (black) and hAPP injected (magenta) conditions (right). L5 SS Ctx sEPSCs show no change in the IEIs (p=0.7372, t=0.3450, df=15; two-tailed unpaired t-test). See Extended Data Fig. 13 for mEPSC data. **Bottom:** Cumulative distribution curve for spontaneous IPSCs in the SS Ctx showing the relationship of relative frequency of events to the inter-event interval (left). Quantified averages of IEIs are displayed for each cell as individual data points and compared between Ctrl (black) and hAPP injected (magenta) conditions (right). L5 SS Ctx sIPSCs show no change in the IEIs (p=0.0812, t=1.890, df=15; two-tailed unpaired t-test). See Extended Data Fig. 13 for mIPSC data.

### hAPP-induced network dysfunction is attenuated following PV interneuron activation *in vivo*

To understand how short-term expression of hAPP affected network behavior *in vivo*, we next performed local field potential (LFP) recordings in the LEC of lightly anesthetized, head-fixed mice 3 weeks after injection of hAPP or control (tdTom) virus (Fig. 6a-c). As predicted by our *ex vivo* studies, we found a broad increase in activity across several frequency domains, similar to previous findings in the LEC of a sporadic AD model^15^. Notably, the delta, theta, and low gamma frequency domains showed significant increases in power (Fig 6e) indicating increases in neuronal activity^33^ (Extended Data Fig. 14b). Despite following a similar trend, frequency domains above 80 Hz (i.e., high Gamma and beyond) were not significantly altered (Fig. 6e, Extended Data Fig. 14b). Together with our mechanistic evaluations earlier, this suggests that hAPP-induced PV interneuron dysfunction induces network hyperexcitability *in vivo*.

**Figure 6.**
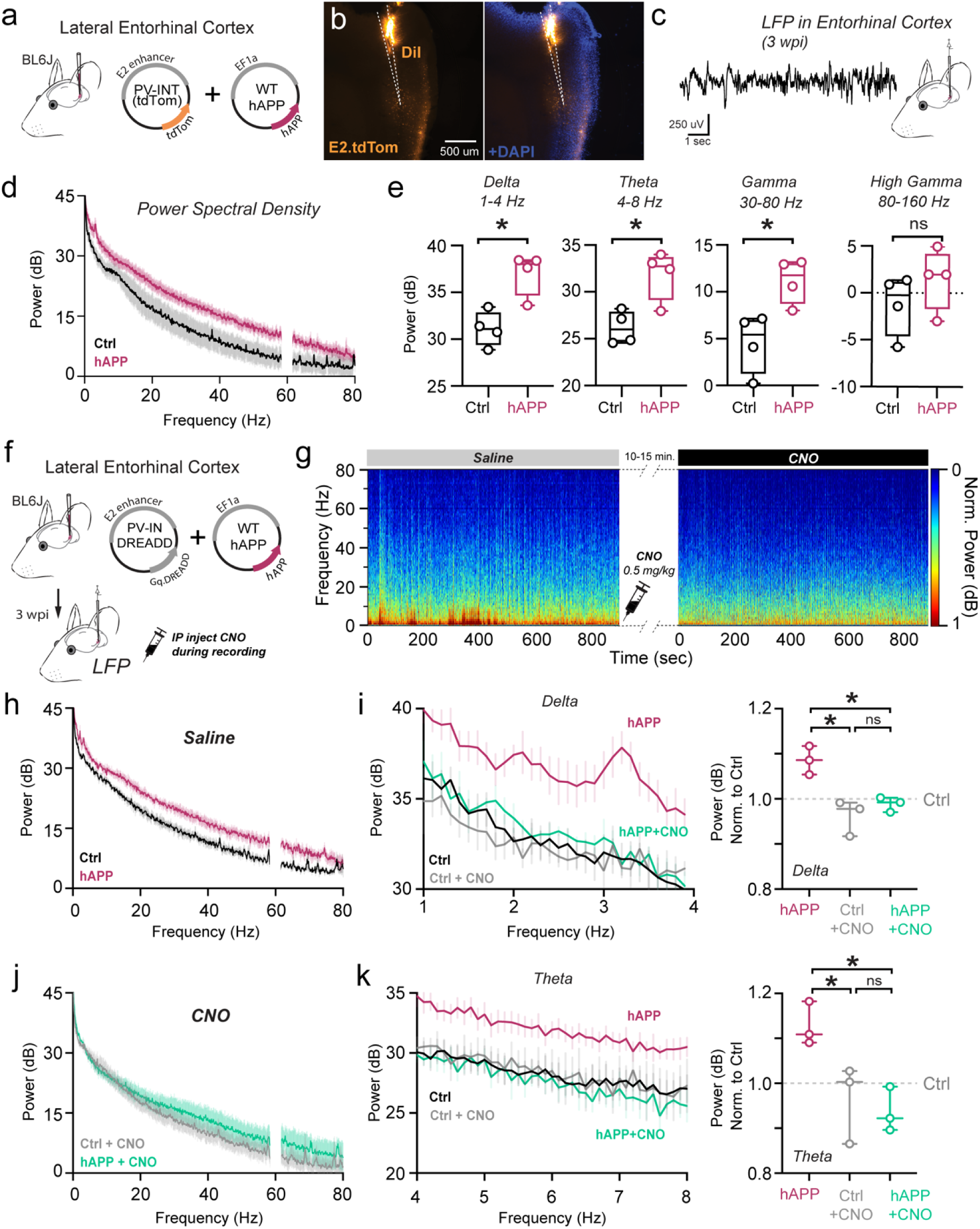
hAPP-induced hyperexcitability observed in vivo can be attenuated by enhancement of PV interneuron excitability. **a.** Graphical summary of AAV.EF1a.hAPP (or for Ctrl, E2.tdTom) stereotactic injection in the Lateral Entorhinal Cortex. **b.** 3 weeks post-injection, mice were anesthetized for *in vivo* glass electrode LFP recordings. (b) is a representative slice at 10x magnification from the LEC where DiI coated electrode recorded from a patch of E2.tdTom+ expressing cells. **c.** Raw LFP signal from *in vivo* glass electrode recordings. **d.** Power Spectral Density [PSD] analysis (dark lines: mean, light lines: ±SEM) comparing Ctrl (E2.tdTom+saline injected) to hAPP (E2.tdTom+hAPP injected). **e.** Power derived from PSD analysis averaged for each mouse in the Delta (1-4 Hz), Theta (4-12 Hz), Gamma (30-80 Hz), and High Gamma (80-160 Hz) frequency ranges and compared between Ctrl and hAPP. Delta (Ctrl: 31.11± 0.94, Min. 28.86, Max. 33.43, Range. 4.57; hAPP: 37.02 ± 1.15, Min. 33.62, Max. 38.43, Range. 4.81; p=0.0073, t=3.98, df=6); Theta (Ctrl: 26.20± 0.88, Min. 24.57, Max. 28.22, Range. 3.65; hAPP: 31.86 ± 1.34, Min. 27.97, Max. 33.98, Range. 6.01; p=0.012, t=3.54, df=6), Gamma (Ctrl: 4.56± 1.59, Min. 0.24, Max. 7.13, Range. 6.89; hAPP: 11.17 ± 1.20, Min. 8.01, Max. 13.15, Range. 5.14; p=0.016, t=3.32, df=6), High Gamma (Ctrl: −1.22± 1.63, Min. −5.76, Max. 1.35, Range. 7.11; hAPP: 1.49 ± 1.65, Min. −2.98, Max. 4.97, Range. 7.96; p=0.2864, t=1.170, df=6). Individual data points and box plots are displayed. Statistical significance is denoted as *p<0.05, as determined by two-tailed unpaired t-test. **f.** Graphical summary of AAV.E2.Gq.DREADD with AAV.EF1a.hAPP (or for Ctrl DREADD, E2.Gq.DREADD + E2.tdTom) stereotactic injection in the Lateral Entorhinal Cortex. 3 weeks post-injection, mice were anesthetized for *in vivo* glass electrode LFP recordings with administration of saline or 0.5 mg/kg administration of Clozapine-n-oxide [CNO] during the recording with a wait time after administration of 10-15 minutes prior to recording. **g.** Spectrogram heatmap of LFP power (normalized) from the recording of one mouse after intraperitoneal saline injection (left, 15 min.) and intraperitoneal CNO injection (right, 15 min.). **h.** Power Spectral Density analysis (dark lines: mean, light lines: ±SEM) comparing Ctrl DREADD (E2.Gq.DREADD + E2.tdTom + saline) to hAPP DREADD (E2.Gq.DREADD + E2.tdTom + hAPP). **i. [Left]** Power Spectral Density analysis only for the Delta frequency range (1-4 Hz) (dark lines: mean, light lines: ±SEM) comparing Ctrl DREADD (black), Ctrl DREADD after CNO (Gray), hAPP DREADD (maroon), and hAPP DREADD after CNO (green). **[Right]** Power [derived from PSD analysis] averaged for each mouse in the Delta frequency range. (hAPP to Ctrl DREADD +CNO: p= 0.0063, hAPP DREADD to hAPP DREADD +CNO: p= 0.0193, Ctrl DREADD + CNO to hAPP DREADD + CNO: p= 0.5758, df=7, One-Way ANOVA with Multiple Comparisons). **j.** Power Spectral Density analysis (dark lines: mean, light lines: ±SEM) comparing Ctrl DREADD after 0.5 mg/kg Clozapine-n-oxide [CNO] injection (E2.Gq.DREADD + E2.tdTom + saline) to hAPP DREADD after 0.5 mg/kg CNO injection (E2.Gq.DREADD + E2.tdTom + hAPP). **k. [Left]** Power Spectral Density analysis only for the Theta frequency range (4-10 Hz) (dark lines: mean, light lines: ±SEM) comparing Ctrl DREADD (black), Ctrl DREADD after CNO (Gray), hAPP DREADD (maroon), and hAPP DREADD after CNO (green). **[Right]** Power [derived from PSD analysis] averaged for each mouse in the Theta frequency range. (hAPP to Ctrl DREADD +CNO: p= 0.0493, hAPP DREADD to hAPP DREADD +CNO: p= 0.0261, Ctrl DREADD + CNO to hAPP DREADD + CNO: p= 0.8593, df=7, One-Way ANOVA with Multiple Comparisons). Also see Extended Data Figure 14 for related datasets.

We next asked whether ‘real-time’ enhancement of PV interneuron excitability could ameliorate hAPP-induced network hyperexcitability *in vivo*. To accomplish this, PV interneurons in the LEC were activated chemogenetically using the E2 PV-specific targeting enhancer^18^ to express hM3Dq.DREADD (Gq.DREADD). For DREADD experiments, groups are denoted as either Ctrl (Gq.DREADD/no hAPP) or hAPP (Gq.DREADD + hAPP) (Fig. 6f,h,j). Remarkably, chemogenetic activation of PV interneurons with low-dose CNO (0.5mg/kg) could rapidly restore the hyperexcitable LFP in hAPP-expressing mice to Ctrl levels (Fig. 6g,h) most notably in the delta (Fig. 6i) and theta (Fig. 6k). Perhaps surprisingly, 0.5mg/kg CNO did not significantly reduce LFP power in Ctrl (no hAPP) animals with respect to saline. Importantly, DREADD expression itself or CNO administration without DREADD expression did not alter LFP power in either the Ctrl or hAPP animal groups (Extended Data Fig. 14c,d). Thus chemogenetic enhancement of PV interneurons in the LEC was sufficient to completely rescue the hAPP-induced hyperexcitability *in vivo*.

### hTau co-expression with hAPP quells LEC hyperexcitability at the cost of increased pathological tau species

Beyond hyperexcitability, the LEC is also the first cortical region to develop tau pathology^1,34–37^. Although Alzheimer’s is characterized by early hAPP/Aβ and later Tau pathology, respectively, the relationship between hAPP, hyperexcitability, and Tau remains unclear. It has previously been established that artificially increasing neuronal activity can accelerate tau pathology^38–40^. However, long-term transgene expression of human Tau (hTau) may act to dampen circuit excitability^16,41,42^ (but see^43^). Thus, we sought to assess the interplay of hAPP-induced circuit hyperexcitability and hTau expression in the LEC. To achieve this, we packaged full-length wild-type human Tau (hTau) into a separate AAV to induce Tau expression locally in the entorhinal cortex. Spontaneous post-synaptic currents were then recorded from LII principal cells, 3 weeks after hAPP alone, hTau alone, or hAPP + hTau co-injection (Fig. 7a). With hAPP, we again observed an elevated E:I frequency ratio (sEPSC frequency/sIPSC frequency, normalized to the Control dataset) as described earlier (Fig. 7b). We hypothesized that hTau would result in a reduced E:I ratio with respect to the control baseline. Although the E:I ratio with hTau alone was less than hAPP alone, E:I balance surprisingly remained unchanged with respect to the Control (Fig. 7b). However, hAPP + hTau resulted in an intermediate effect, which abolished the hyperexcitable phenotype seen with hAPP alone (Fig. 7b). These results agree with a homeostatic role for Tau in maintaining circuit excitability. Beyond synaptic event frequencies, all other spontaneous event properties (i.e., amplitude) were statistically similar between all groups (Extended Data Fig. 15a,b).

**Figure 7.**
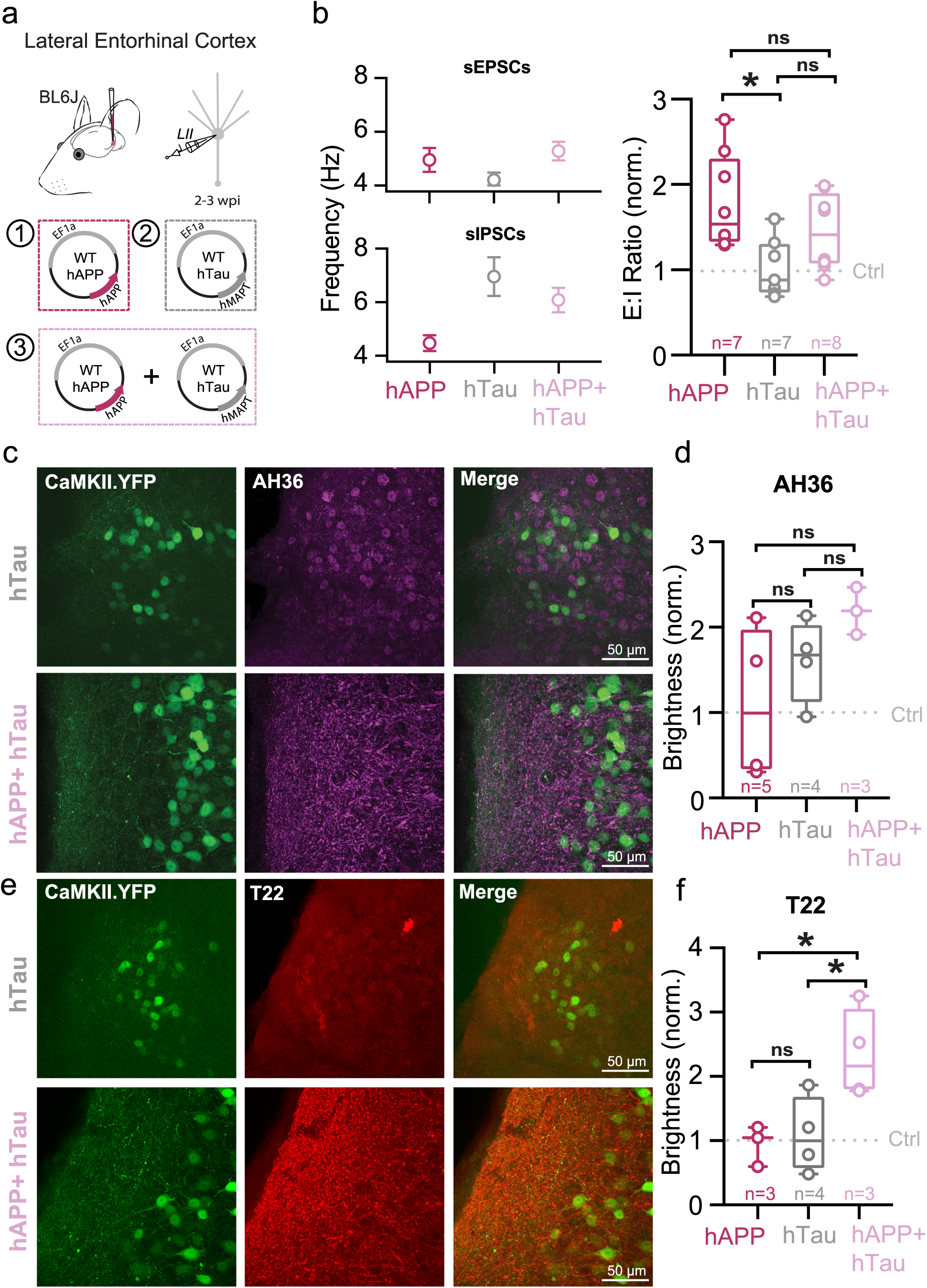
hTau co-expression with hAPP quells hyperexcitability but increases pathological tau. **a.** Graphical summary of AAV.EF1a.hAPP, AAV.EF1a.MAPT (hTau), or co-injected AAV.EF1a.hAPP with AAV.EF1a.MAPT stereotactic injection in the Lateral Entorhinal Cortex. Excitatory cells were targeted for whole-cell voltage-clamp recordings. **b.** Spontaneous events obtained by holding cell voltage at −70 mV (excitatory post-synaptic currents, EPSCs [top]) and 0 mV (inhibitory post-synaptic currents, IPSCs [bottom]), interleaved. Quantified averages of event frequency are displayed for each cell normalized to Ctrl values as a ratio of EPSC Frequency to IPSC frequency and compared between hAPP injected (magenta), hTau injected (gray) and hAPP + hTau co-injected (pink) conditions. L2 LEC injected with hAPP showed a significantly elevated E:I ratio compared to hTau injected (p=0.0136, df=20). hAPP and hTau co-injected E:I ratio was not significantly different from hAPP injected (p=0.3323, df=20) or hTau injected (p=0.2175, df=20). For all summary graphs, data are expressed as mean (± SEM). Statistical significance is denoted as *=p<0.05, as determined by an Ordinary one-way ANOVA with Multiple comparisons. **c,e.** IHC representative images at 60x magnification for hTau (top) or hAPP+hTau (bottom) injected mice (for Ctrl or hAPP injected, see Extended Data Fig. 15) with staining for either AH36 (c) or T22 (e). **d.** hAPP, hTau, and hAPP+hTau were analyzed for AH36 brightness in the first 100 um of every slice. AH36 brightness was normalized to CaMKII.eYFP brightness to control for any potential variability in viral expression. All groups were then normalized to the Ctrl injected condition. hAPP+hTau showed the highest level of AH36 brightness, although it was not significant over hAPP (p=0.1267) or hTau (p=0.4900) (df=8, One-Way ANOVA with Multiple Comparisons). hAPP and hTau were also not significantly different (p=0.5328). **e.** hAPP, hTau, and hAPP+hTau were analyzed for T22 brightness in the first 100 um of every slice. AH36 brightness was normalized to CaMKII.eYFP brightness to control for any potential variability in viral expression. All groups were then normalized to the Ctrl injected condition. hAPP+hTau showed a significantly higher level of T22 brightness, above both hAPP (p=0.0350) and hTau (p=0.0.0389) (df=8, One-Way ANOVA with Multiple Comparisons). hAPP and hTau were not significantly different (p=0.9526).

We next assessed whether the moderating effect of hTau on circuit activity came at the cost of increased pathology, using antibodies for pSer202/pThr205 phosphorylated tau (AH36) or oligomeric tau (T22). Both Control and hAPP-injected conditions showed low levels of AH36 positivity, likely due to endogenous labeling of murine tau (Extended Data Fig. 15c,d). While both hTau and hAPP+hTau induced high levels of AH36-positive staining (Fig. 7c,d), it appeared that hTau alone injected mice had mostly somatically located staining. In contrast, hAPP+hTau co-injected mice displayed disperse staining (Fig. 7c) suggesting an interaction with hAPP which promotes extracellular transport of pathological Tau. Oligomeric tau (T22) (Fig. 7e), which has recently been shown in human tissue as a tau species that may spread transsynaptically from axons to other regions^44^, displayed a surprisingly robust increase, but only when hAPP+hTau were co-expressed (Fig. 7e,f; Extended Data Fig. 15e,f). Thus, it appears that co-expression of Tau could help restore APP-induced circuit hyperexcitability. However, a consequence of their co-expression appears to be the amplification and translocation of known pathological tau species.

## Discussion

Here we demonstrate that PV interneurons within the LEC are biophysically distinct from other neocortical PV interneurons. Furthermore, differences in the native-state proteomes of PV interneurons from the LEC and SS Ctx regions were marked. Although the WT PV firing frequency in our LEC recordings is consistent with previous obervations^17^, the striking biophysical differences (i.e., AP waveform) with respect to PV cells in other cortical regions had not been systematically evaluated. Interestingly, these LEC PV interneurons do resemble a previously observed PV+ interneuron in other regions, such as the ‘quasi fast-spiking interneurons’ of the subiculum^45^, the ‘fast-spiking-like cells’ of the striatum^46^, and the ‘non-fast-spiking interneurons’ of the CA1^47^. However, where these cells represent a small subset of the PV+ interneurons of these regions (∼20% in CA1)^47^, the majority of our recorded PV+ interneurons displayed this low-firing phenotype. Whether the baseline low-firing frequency of PV+ interneurons in the LEC confers vulnerability to hAPP-induced pathophysiology remains unclear. In addition to their different intrinsic features, PV interneurons of the LEC displayed a multitude of differentially expressed proteins in comparison to SS Ctx PV interneurons. Interestingly, compared to SS, we found that PV interneurons residing in the LEC were significantly enriched in proteins associated with cognitive resilience in humans. However, many of these LEC PV-IN pro-resilience proteins were altered in the entorhinal cortex of AD patients. This suggests a regional and cell-type-specific susceptibility to the progression of AD pathophysiology. Although a comparison of PV-CIBOP regional proteomes with bulk human brain proteomes gives further insight into potential cell-type-specific alterations in phases of AD, it is still limited by the inability to verify PV interneuron-specificity of observed changes in human brain. As we only recently established the first, to our knowledge, PV interneuron-specific proteome^21^, we look forward to the advancement of techniques to come in order to complete such a level of analysis in human tissue. Future studies at the single-cell level in humans with early-stage AD will be necessary to confirm this assertion.

Recent work shows that APP expression moves outside of normal homeostatic levels in models of late-onset AD risk alleles^48,49^. The ratio of different APP isoforms also shifts in human AD, from mainly APP 695 to increasing levels of APP 770 and 751^26,27^. These longer isoforms show increased expression following aging-related processes (e.g., after reproductive hormonal production decline^50^, hypercholesterolemia^51^, and atherosclerosis^52^), all of which are also associated with increased AD risk^53–56^. Thus, here we induced adult-onset expression of hAPP 770 to model these phenomena. Adult-onset expression of hAPP allowed us to avoid any alterations to neurodevelopment which may arise with expression of a transgene early in development, as many mouse models of AD exhibit. However, we acknowledge that this model does not encapsulate all alterations that may arise throughout aging and early Alzheimer’s Disease. Thus, further studies must be conducted to assess these mechanisms in aging mice. At this time point, we found that shortly after hAPP expression (2-3 weeks), LEC PV interneuron firing became severely disrupted. Although viral hAPP expression was robust in both excitatory and inhibitory cells due to the generic promoter used to express hAPP in our experiments, we observed no alteration to the intrinsic excitability of excitatory cells in LEC LII. Despite this, there was a significant disruption in the E:I balance in LEC slices. Thus, we propose basal network hyperexcitability observed arises as a result of decreased PV-interneuron firing, resulting in a loss of tonic inhibition and thus, increased firing propensity of surrounding excitatory neurons. Notably, our hAPP condition displayed a large peak in LFP power in the delta frequency domain (1-4 Hz) *in vivo*, which was returned to baseline upon rescue of PV interneuron excitability. This delta peak may represent increased low-frequency excitatory neuron synchronicity. We also observed a broadband increase in most other frequency ranges in the hAPP condition, including gamma power, which has been previously described in AD patients^57^ and in amyloid pathology mouse models^13^.

Of the 26 amino acids differentiating the hAPP and mAPP proteins, only 3 are situated within the amyloid-beta region. Of note, of the ‘wild-type’ versions of newly designed hAPP knock-in mouse models^58,59^ now in wide use, only the 3 amino acids within the amyloid region are humanized. It has been shown that expression of WT hAPP can result in a substantial increase in amyloid-beta^60^, suggesting that PV interneuron hypofunction in the LEC phenotype results directly from Aβ accumulation. Thus, we investigated if humanizing only the 3 amino acids could recapitulate our findings of hAPP-induced impaired PV interneuron physiology. Interestingly, the mAPP/hAβ Chimera did result in impaired PV-interneuron firing, but did not replicate the more drastic alterations seen with full-length hAPP expression. This suggests amyloid-beta is not the sole cause for early phase interneuron dysfunction, and may suggest a role for either full-length APP or its other cleavage products in this stage of the neurodegenerative cascade. Alternatively, the full-length hAPP may result in enhanced Aβ production with respect to the mAPP/Aβ chimera.

GABAergic interneurons require homeostatic APP levels for proper physiological function and circuit activity control^61^. Furthermore, APP^62^, as well its cleavage proteins^63–65^ and products^66,67^, can modulate neuronal biophysics and alter the expression of ion channels, many of which are essential for maintaining the ‘fast-spiking’ phenotype of PV interneurons. Modifications to Na_v_1 or K_v_3 channel availability in different constitutive hAPP-expressing mice have recently been linked to reduced PV excitability^7,13^. Although short-term full-length hAPP expression in this study could significantly reduce PV firing, we observed no outward biophysical indicators implicating changes to either Na_v_1 or K_v_3 availability which could underlie altered PV firing in the LEC. However, this likely requires further evaluation of specific ion channel subtypes.

Although firing of PV interneurons in SS cortex were unaffected with short-term hAPP expression, we did note a change in AP threshold, potentially attributable to alterations in Na_v_ channels. It is likely that if expressed longer, hAPP expression in the SS Ctx may result in impaired PV interneuron excitability^7,13^. Thus, alternative biophysical mechanisms are likely responsible for our observations following short-term hAPP expression in adult mice. Notably, we observed a substantial decrease in input resistance in LEC PV cells expressing hAPP. This could be due to enhanced availability of leak channels or potentially low-voltage activating K^+^ conductances, such as KCNQ (K_v_7), which have been shown to be regulated by APP cleaving proteins^63^ and cleavage products^66,68,69^. Interestingly, the Gq-coupled hM3Dq DREADD system increases cell excitability specifically by increasing input resistance in cells. Of note, here we show that recovery of reduced input resistance in PV interneurons through specific activation of Gq.DREADDs was sufficient to attenuate hAPP-induced hyperexcitability in the LEC *in vivo*.

The LEC is also the first cortical region to develop tau pathology^1,34–37^. Yet, the relationship between hAPP, hyperexcitability, and Tau remains unclear. It has previously been established that artificially increasing neuronal activity can accelerate tau pathology^38–40^. However, the expression of hTau has been suggested to strongly dampen circuit excitability^16,41,42^ (but see^43^). Here we observed that hTau co-expressed with hAPP results in an intermediate circuit excitability level when compared to hAPP or hTau injected alone. Synaptic spread of tau has been shown from the entorhinal cortex to other brain regions^70,71^, and most recently this effect has been suggested to occur in human brain via the oligomeric tau species (T22+)^44^. Remarkably, although hTau co-injection with hAPP somewhat normalized circuit excitability, it also resulted in a significant increase and apparent extracellular translocation of phospho-tau and oligomeric tau species.

The LEC is the first cortical region to undergo end-stage cellular neurodegeneration^1^ in AD, specifically, Layer II^2^ excitatory cells^4^. Conversely, one of the earliest pathophysiological alterations seen in both humans with AD, and in mouse models of early- and late-onset AD pathology^9,15,16^ is altered local circuit excitability^3,72,73^. In agreement with our mechanistic cellular and *in vivo* findings here, hyperactivity has been shown to preferentially emerge in the LEC region^3^. As with others, our slice electrophysiology assessments of circuit excitability aligned with our *in vivo* findings as good predictors of hyperexcitability in AD pathology studies^7,14,74^. Our study suggests that hAPP-induced hyperexcitability in the LEC arises not from alterations in the intrinsic or synaptic properties of AD-vulnerable LII excitatory cells, but rather from an initial alteration in intrinsic excitability of surrounding interneurons. The fact that short-term hAPP expression in SS cortex caused {no changes in PV firing or overall basal circuit excitability supports this notion. Circuit hyperexcitability is likely an influential factor in the early neurodegenerative cascade^7575^, as it has been shown to exacerbate release of amyloid-beta^76^, and also promotes tau pathology and subsequent trans-synaptic tau spread^39^ which ultimately tracks with spine synapse degeneration^14^ and cell death^77^. Ultimately, regions that first undergo hyperexcitability may also be among the earliest to display these pathological markers as the disease progresses^39,75^. Thus, PV interneurons of the LEC potentially represent a prime target for therapeutic intervention in the early stages of Alzheimer’s Disease.

## Supporting information

Supplemental Data Proteomics 1

Supplemental Data Proteomics 2

Supplemental Data Proteomics 3

Supplemental Data Proteomics 4

## Methods

### Acute slice preparation

All animal procedures were approved by the Emory University IACUC. Acute slices from SS cortex and LEC were prepared from a mixture of C57Bl/6J and PV-Cre mice (8-12 weeks old), evenly dispersed between test and control groups. Male and female mice were used for all experiments with data collected from ≥ 3 mice per experimental condition. Mice were first anesthetized and then killed by decapitation. The brain was then immediately removed by dissection in ice-cold cutting solution (in mM) 87 NaCl, 25 NaHO3, 2.5 KCl, 1.25 NaH2PO4, 7 MgCl2, 0.5 CaCl2, 10 glucose, and 7 sucrose. Brain slices (250 μm) were sectioned in the sagittal plane using a vibrating blade microtome (VT1200S, Leica Biosystems) in the same solution. Slices were transferred to an incubation chamber and maintained at 34°C for ∼30 min and then at 23-24°C thereafter. During whole-cell recordings, slices were continuously perfused with (in mM) 128 NaCl, 26.2 NaHO3, 2.5 KCl, 1 NaH2PO4, 1.5 CaCl2, 1.5MgCl2 and 11 glucose, maintained at 30.0±0.5°C. All solutions were equilibrated and maintained with carbogen gas (95% O2/5% CO2) throughout.

### Slice Electrophysiology

PV interneurons and excitatory cells were targeted for somatic whole-cell recording in layer 5 region of somatosensory cortex or layer 2 lateral entorhinal cortex by combining gradient-contrast video-microscopy with epifluorescent illumination on custom-built or commercial (Olympus) upright microscopes. Electrophysiological recordings were acquired using Multiclamp 700B amplifiers (Molecular Devices). Signals were filtered at 10 kHz and sampled at 50 kHz with the Digidata 1440B digitizer (Molecular Devices). For whole cell recordings, borosilicate patch pipettes were filled with an intracellular solution containing (in mM) 124 potassium gluconate, 2 KCl, 9 HEPES, 4 MgCl2, 4 NaATP, 3 L-Ascorbic Acid and 0.5 NaGTP. For experiments recording spontaneous and miniature events, an intracellular solution containing (in mM) 120 CsMeSO4, 10 HEPES, 5 TEA.Cl, 4 Na2ATP, 0.5 Na2GTP, 2 MgCl2, 10 L-Ascorbic Acid, and 3 Qx314. To obtain miniature events, slices were perfused with 1 μM TTX for 10 minutes prior to recording (at least 10 mL run). The same protocol was used for spontaneous and miniature events. In patch experiments using exogenously applied amyloid, slices were incubated in an equal mixture of 0.25 μM Aβ_1-40_ and Aβ_1-42_, as described previously^10^. LEC slices were incubated with this solution for 3h before being transferred to standard extracellular solution for patch recordings. Pipette capacitance was neutralized in all recordings and electrode series resistance compensated using bridge balance in current clamp. Liquid junction potentials were uncorrected. Recordings had a series resistance < 20 MΩ. Membrane potentials were maintained near −70 mV during current clamp recordings using constant current bias. Action potential trains were initiated by somatic current injection (300 ms) normalized to the cellular capacitance in each recording measured immediately in voltage clamp after breakthrough^78^. For quantification of individual AP parameters, the 1^st^ AP in a spike train at was analyzed at 12 pA/pF for all cells. Passive properties were determined by averaging the responses of several 100 ms long, −20pA steps during each recording. Spontaneous and miniature events were recorded at a holding voltage of −70 and 0 mV, one second each, interleaved for 3-5 minutes. Statistical analyses of spontaneous and miniature events using cell averages, to avoid bias in favor of cells which may receive more inputs or less^74,79^. For regional comparisons of PV interneurons, combined controls from datasets in each region were used. Event detection was carried out using Clampfit (Molecular Devices) using a template matching algorithm and were curated manually with a 4 kHz low-pass filter.

### In vivo Electrophysiology

Electrophysiological recordings (Local Field Potential) were performed in lightly anesthetized animals (1% v/v isoflurane). Pipettes (1–3 MΩ) with a long shank (∼2 cm) were pulled from borosilicate glass capillaries (1.65mm O.D, #PG52615-4, Sutter instruments Co., USA) using a vertical puller (Narishige PC-100), coated with Sylgard (DOW), baked in a convection oven at 145 °F for 30 min, and allowed to cool prior to use. The reference electrode (silver chloride) was implanted in the contralateral cerebellum. For recordings, pipettes were filled with sterile saline, coated with DiI, slightly trimmed, and placed onto the silver chloride wire. Positive pressure (52-53.9 mBar) was applied to the pipettes during entry into the cortex and then reduced to <1.0 mBar prior to recording. Recordings were conducted at the same coordinates as the injection site in the LEC (X = ± 3.39; Y = −4.52; α = 0°, Z= −2.0). Electrophysiological recordings were acquired using Multiclamp 700B amplifiers (Molecular Devices). Signals were filtered at 4 kHz (Bessel) and sampled at 10 kHz with the Digidata 1440B digitizer (Molecular Devices). For each session, 3-5 5 minute recording sessions were completed. Each animal was first injected intraperitoneally with sterile saline, and after a 15 minute wait period were recorded. Subsequently, each animal was injected with 0.5 mg/kg Clozapine N-oxide dihydrochloride (#6329, TOCRIS) reconstituted in sterile saline. Data was analyzed offline. 30 minutes of recording (total) were analyzed for each mouse: 15 minutes of saline IP injected condition and 15 minutes of the CNO injected condition. LFP recordings were subjected to a fast-Fourier transformation (FFT) and converted to Power in decibels using custom code to achieve a power spectral density analysis. Data points for each frequency range (Delta-High Frequency Oscillations) were averaged for each mouse in each condition to compare for statistical analysis. Statistical analysis was completed using GraphPad Prism. Confirmation of electrode location was completed post-hoc with imaging of LEC sagittal slices on the Nikon Eclipse Ti-E microscope.

### Intracranial viral injections

5-9 week old mice were injected with AAV(PHP.eB).E2.tdTom with saline or AAV(PHP.eB).EF1a.hAPP (0.3 μL total, 1:1) in the SBFI vibrissal region of cortex or the Lateral Entorhinal Cortex. For murine APP experiments AAV(PHP.eB).EF1a.hAPP was replaced with AAV(PHP.eB).EF1a.mAPP. For murine APP/hAB chimera experiments, AAV(PHP.eB).EF1a.mAPP/hAB was used. For tau experiments, the four conditions were: Ctrl (CaMKII.eYFP: saline 1:2), hAPP (EF1a.hAPP: CaMKII.eYFP: saline, 1:1:1), hTau (AAV(PHP.eB).EF1a.hMAPT: CaMKII.eYFP: saline, 1:1:1), and hAPP+hTau (EF1a.hAPP: Ef1a.hMAPT: CaMKII.eYFP, 1:1:1). For DREADD experiments, animals were injected with AAV.E2.Gq.DREADD and AAV(PHP.eB).E2.tdTom and either AAV(PHP.eB).EF1a.hAPP or sterile saline (1:1:1). When performing intracranial viral injections, mice were head-fixed in a stereotactic platform (David Kopf Instruments) using ear bars, while under isoflurane anesthesia (1.5 - 2.0%). Thermoregulation was provided by a heating plate using a rectal thermocouple for biofeedback, thus maintaining core body temperature near 37°C. Bupivacaine was subcutaneously injected into the scalp to induce local anesthesia. A small incision was opened 5-10 minutes thereafter and a craniotomy was cut in the skull (< 0.5 μm in diameter) to allow access for the glass microinjection pipette. Coordinates (in mm from Bregma) for microinjection in the SS Cortex were X = ± 0.85; Y = −2.5; α = 0°; Z = −0.85, coordinates for the LEC were X = ± 3.39; Y = −4.52; α = 0°, Z= −2.4, −1.5. Viral solution (titer 1×10^09^ to 1×10^13^ vg/mL) was injected slowly (∼0.02 μL min-1) by using a Picospritzer (0.3 μL total). After ejection of virus, the micropipette was held in place (5 min) before withdrawal. The scalp was closed with surgical sutures and Vetbond (3M) tissue adhesive and the animal was allowed to recover under analgesia provided by injection of carprofen and buprenorphine SR. After allowing for onset of expression, animals were sacrificed acute slices were harvested.

### Retro-orbital (RO) injection

Male and female mice were given AAV retro-orbital injections as previously described^80^. Mice were anesthetized with 1.8-2% isoflurane. AAV(PHP.eB).E2.Cre.2A.GFP virus was titrated to 2.4×10^11^ vector genomes total was accompanied by AAV(PHP.eB).Flex.tdTom titrated to 3.1×10^11^ and injected in TurboID+ mice^20^ to label PV interneurons throughout cortex. Titrated virus was injected into the retro-orbital sinus of the left eye with a 31G x 5/16 TW needle on a 3/10 mL insulin syringe. Mice were kept on a heating pad for the duration of the procedure until recovery and then returned to their home cage. After 3 weeks post-injection, mice were provided with biotin water continuously. Biotin water was administered for 2 weeks until acute slice sample collection (total of 5 weeks post-RO injection).

### CIBOP studies

PV-CIBOP studies were performed by single retro-orbital injections of AAV(AAV(PHP.eB).E2.Cre.2A.GFP) as described above to Rosa26TurboID/wt mice and WT littermate animals at 7 weeks of age, as previously described^21^. Control groups in PV-CIBOP studies also received AAV E2.Cre injections for fair comparisons. After 3 weeks to allow Cre-mediated recombination and TurboID expression, biotinylation (37.5 mg/L in drinking water) was performed for 2 weeks^20^. After 5 weeks (total), acute slices were acquired as described above, with subsequent microdissection of the SS Ctx or the LEC.

### Tissue processing for protein-based analysis, including Western Blot (WB)

Tissue processing for proteomic studies, including Mass Spectrometry (MS), were performed similarly to previous CIBOP studies^20,21^. Frozen brain tissues (whole brain homogenate excluding cerebellum for WB and microdissected cortical regions for Fig 1) either intact or dissected cortex, was weighed and added to 1.5mL Rino tubes (Next Advance) containing stainless-steel beads (0.9-2mm in diameter) and six volumes of the tissue weight in urea lysis buffer (8 M urea, 10 mM Tris, 100 mM NaH2PO4, pH 8.5) containing 1X HALT protease inhibitor cocktail without EDTA (78425, ThermoFisher). Tissues were homogenized in a Bullet Blender (Next Advance) twice for 5 min cycles at 4 °C. Tissue were further sonicated consisting of 5 seconds of active sonication at 20% amplitude with 5 seconds incubation periods on ice. Homogenates were let sit for 10 minutes on ice and then centrifuged for 5 min at 12,000 RPM and the supernatants were transferred to a new tube. Protein concentration was determined by BCA assay using Pierce™ BCA Protein Assay Kit (23225, Thermofisher scientific). For WB analyses, 10μg of protein from brain lysates were used to verify TurboID expression (anti-V5) and biotinylation (streptavidin fluorophore conjugate). Standard WB protocols, as previously published, were followed^20^. All blots were imaged using Odyssey Infrared Imaging System (LI-COR Biosciences) or by ChemiDoc Imaging System (Bio-Rad) and densitometry was performed using ImageJ software.

### Enrichment of biotinylated proteins from CIBOP brain

As per CIBOP protocols previously optimized^20^, biotinylated proteins were captured by streptavidin magnetic beads (88817; Thermofisher Scientific) in 1.5 mL Eppendorf LoBind tubes using 83uL beads per 1mg of protein (here 200 µg of protein with 16.6 µL of beads) in a 500 μL RIPA lysis buffer (RLB)(50 mM Tris, 150 mM NaCl, 0.1% SDS, 0.5% sodium deoxycholate, 1% Triton X-100). In brief, the beads 16.6 µL beads were washed twice with 1 ml of RLB and 200µg of protein were incubated by making up the total volume of the solution up to 500 µl using RPL. After incubation at 4 deg C for 1 h with rotation, beads were serially washed at room temperature (twice with 1 mL RIPA lysis buffer for 8 min, once with 1 mL 1 M KCl for 8 min, once with 1 mL 0.1 M sodium carbonate (Na2CO3) for ∼10 s, once with 1 mL 2 M urea in 10 mM Tris-HCl (pH 8.0) for ∼10 s, and twice with 1 mL RIPA lysis buffer for 8 min), followed by 1 RIPA lysis buffer wash 4 final PBS washes. Finally, after placing the tubes on the magnetic rack, PBS was removed completely, then the beads were further diluted in 100 μl of PBS. The beads were mixed and 10% of this biotinylated protein coated beads were used for quality control studies to verify enrichment of biotinylated proteins (including WB and silver stain of proteins eluted from the beads). Elution of biotinylated protein was performed by heating the beads in 30 μL of 2X protein loading buffer (1610737; BioRad) supplemented with 2 mM biotin + 20 mM dithiothreitol (DTT) at 95°C for 10 min. The remaining 90% of sample were stored at −20°C for western blot or mass spectrometric analysis of biotinylated protein.

### Western blotting and Silver Stain

To confirm protein enrichment on the pulldown samples the protein were eluted from the beads as mentioned above and 1/3^rd^ of the eluted protein was resolved on a 4–12% Bris-Tris gel (Invitrogen: cat# NW04125box) and transferred onto transferred onto iBlot 2 Transfer Stack containing nitrocellulose membrane using the BOLT transfer system. The membranes were washed once with TBS-T (0.1% tween-20) and then blocked with Start Block (37543, Thermofisher Scientific) for 1 h. The membrane was then probed with streptavidin-Alexa-680 diluted in Start Block for 1 h at room temperature. Further, the membranes were washed 3 times and biotinylated proteins were detected on Odyssey Infrared Imaging System (LI-COR Biosciences). In parallel, 2/3^rd^ of the eluted protein samples were resolved on a 4–12% Bris-Tris gel (Invitrogen: cat# NW04125box) and subjected to Silverstein (Pierce™ Silver Stain Kit: Catalog number: 24612) to detect the total amount of protein in each lane according to the manufactured protocol.

### Protein digestion, MS, protein identification and quantification

On-bead digestion of proteins (including reduction, alkylation followed by enzymatic digestion by Trypsin and Lys-C) from SA-enriched pulldown samples (1mg protein used as input) and digestion of bulk brain (input) samples (25 μg protein), were performed as previously described with no protocol alterations^20,21^. In brief, after the removal of PBS from remaining 90% of streptavidin beads (10% used for quality control using western blot and silver stain) were resuspended in 90 uL of 50 mM ammonium bicarbonate (NH4HCO3) buffer. Biotinylated proteins were then reduced with 1 mM DTT and further alkylated with 5 mM iodoacetamide (IAA) in the dark for 30 min each on shaker. Proteins were digested overnight with 0.2 μg of lysyl (Lys-C) endopeptidase (127-06621; Wako) at RT on shaker followed by further overnight digestion with 0.4 μg trypsin (90058; ThermoFisher Scientific) at RT on shaker. The resulting peptide solutions were acidified to a final concentration of 1% formic acid (FA) and 0.1% triflouroacetic acid (TFA), desalted with a HLB columns (Cat#186003908; Waters). The resulting protein solution was dried in a vacuum centrifuge (SpeedVac Vacuum Concentrator). Detailed methods for this protocol have been previously published^20^. Lyophilized peptides were resuspended followed by liquid chromatography and MS (Q-Exactive Plus, Thermo, data dependent acquisition mode) as per previously published protocols^20^. As previously published^81–83^, MS raw data files were searched using SEQUEST, integrated into Proteome Discoverer (ThermoFisher Scientific, version 2.5) using the Uniprot 2020 database as reference (91,441 target 37 sequences including V5-TurboID). Raw MS data as well as searched Proteome Discoverer data before and after processing to handle missing values, will be uploaded to the ProteomeXchange Consortium via the PRIDE repository^84^. The false discovery rate (FDR) for peptide spectral matches, proteins, and site decoy fraction were 1 %. Other quantification settings were similar to prior CIBOP studies^20^. Quantitation of proteins was performed using summed peptide abundances given by Proteome Discoverer. We used razor plus unique peptides for protein level quantitation. The Proteome Discoverer output data were uploaded into Perseus (Version 1.6.15) and abundance values were log2 transformed, after which data were filtered so that >50% of samples in a given CIBOP group expected to contain biotinylated proteins, were non-missing values. Protein intensities from SA-enriched pulldown samples (expected to have biotinylated proteins by TurboID) were normalized to sum column intensities prior to comparisons across groups. This was done to account for any variability in level of biotinylation as a result of variable Cre-mediated recombination, TurboID expression and/or biotinylation^20^.

### Analysis of enrichment of cognitive resilience proteins in regional PV interneuron DEPs

We cross-referenced PV-CIBOP regional proteins (N=207, unadjusted p<0.05) with proteins previously found to be associated with cognitive decline in humans (N=55 proteins associated with cognitive slope in the Religious Orders Study and the Rush Memory and Aging Project (ROSMAP), unadjusted p<0.05)^21,22^. In this prior study, cognitive slope was estimated in the ROSMAP longitudinal study of aging, and cognitive slope was correlated with post-mortem protein abundances, measured by quantitative MS. Proteins that were associated with stable cognitive function (positive association with cognitive slope) were identified as protein markers of cognitive resilience (or pro-resilience proteins). Conversely, proteins ante-correlated with cognitive stability were labeled as anti-resilience proteins. 55 resilience-associated proteins in ROSMAP were identified as regional DEPs in our PV-CIBOP study. The median correlation with cognitive slope for SS Ctx and LEC PV-CIBOP DEPs was estimated and compared across the two regions (two-tailed Mann-Whitney U test).

### Integration of PV-CIBOP proteomes with existing human AD proteomic datasets

We cross-referenced our PV-CIBOP regional proteomes with existing human post-mortem proteomes in which the entorhinal cortex (EC), frontal cortex (FC) and other regions were sampled from AD cases (early BRAAK stages I-III, late BRAAK stages IV-VI) and non-AD/non-pathology controls. Proteins were identified as DEPs if they were significantly different comparing AD versus controls within any given region (total of 737 DEPs identified). PV-CIBOP proteins identified by our current study were further assessed for evidence of regional differences within PV interneurons (SS Ctx vs. LEC).

### Assessment of human tau/APP protein-protein interactors among PV-CIBOP regional DEPs

We cross-referenced our PV-CIBOP regional proteomic data with human tau protein interactors, previously identified in a meta-analysis across 12 published tau interactome studies^24^, in which 2,084 human tau interactors were identified, and among these, 261 were high-confidence interactors if they were identified by at least 3 studies. PV-CIBOP proteins in our study that were also found among these 216 human tau interactors, were further analyzed for evidence of regional differences (SS Ctx vs. LEC) in PV interneurons. The proportions of tau interactors and non-tau interactors across SS Ctx-enriched, LEC-enriched and non-regional PV interneuron proteins were compared (Chi square test).

Using a similar approach, we obtained a list of 243 human APP interactors derived from protein-protein-interaction databases (STRING consortium 2023, Version 12.0, https://string-db.org/) restricting the interactome to physical interactors with medium confidence stringency (confidence scores >0.40, including text mining, experiments and databases). We excluded proteomic studies of extracellular amyloid beta plaques as these are less likely to represent interactions relevant to intra-neuronal APP processing.

### RNAscope

RNAscope was performed for confirmation of viral hAPP and mAPP expression, as well as quantification of viral expression in a cell-type-specific manner. RNAscope was performed as instructed by Advanced Cell Diagnostics (ACD). Tissue was prepared from C57Bl/6J mice (8-12 weeks old), evenly dispersed between test and control groups. Male and female mice were used for all experiments with data collected from ≥ 3 mice per experimental condition. Mice were injected with AAV(PHP.eB).E2.tdTom and AAV(PHP.eB).hAPP or AAV(PHP.eB).mAPP (0.3 uL total, 1:1) in one hemisphere and injected contralaterally with AAV(PHP.eB).E2.tdTom and saline (0.3 uL total, 1:1), n=3 each. Hemispheres were randomized. After 2-3 weeks, mice were first anesthetized and then killed by decapitation. The brain was then immediately removed and flash frozen in isopentane on dry ice. Samples were kept at −-80°C prior to sectioning. Tissue was sectioned on a cryostat at 16 μm thickness, mounted onto Superfrost Plus Slides, and stored at - 80°C until use. Samples were fixed and dehydrated according to the RNAscope kit manufacturer (ACDBio) standard protocol. In brief, frozen slides containing tissue sections were immediately dipped in pre-chilled 4% paraformaldehyde (PFA) for 15 minutes at room temperature. After fixation, slides were briefly rinsed with 1× phosphate buffered saline (PBS) two times to remove excess fixative. Tissue sections were then dehydrated in a series of ethanol solution 50%, 70% and 100% for 5 min at room temperature. After ethanol washes the slides were transferred into a fresh 100% ethanol solution to sit overnight at −20°C. The following day, slides were taken out of the ethanol solution, air dried for 5 minutes, and hydrophobic barriers were drawn around each section. The remainder of the RNAscope assay was then performed following the manufacturer’s protocol, multiplexing two different probe groups on two different sections from each animal: 1) human APP (catalog #418321), mouse APP (catalog #519001-C2), and CaMKIIα (catalog #445231-C3), or 2) human APP, mouse APP, and Pvalb (catalog #421931-C3).

### Immunohistochemistry

Acute slices were acquired as previously described at 100 μm thickness. Immediately following collection from the vibratome, free-floating sections were placed in 4% paraformaldehyde for fixation at room temperature for 1-2 hours. Sections were then washed three times in 1x Tris Buffered Saline (1xTBS) for 10 minutes. To block nonspecific binding, the sections were then incubated with 5% goat serum (in 1xTBS) for 1 hour at room temperature. Sections were then incubated overnight at 4°C on a shaker plate in the primary antibody solution, which contained 0.2% Triton X-100, 1% goat serum, and a 1:1000 dilution of either the AH36 antibody (StressMarq Sciences, #SMC-601) or T22 antibody (Millipore Sigma, #ABN454) in 1xTBS. The next day, sections were washed three times for 10 minutes in 1xTBS before incubation with the secondary antibody (Alexa Fluor^TM^ 647 at 1:1000; ThermoFisher Scientific, #A-21245) in 1xTBS for 1 hour at room temperature on a shaker plate. From the point of secondary antibody incubation, sections were protected from light using aluminum foil. Following secondary antibody incubation, sections were washed again three times for 10 minutes in 1xTBS, mounted on slides (Fisher Scientific, #1255015), and coverslipped with Fluoromount containing DAPI (ThermoFisher Scientific, #00-4959-52). Slides were then imaged on a Nikon C2 laser-scanning confocal system with an inverted Nikon ECLIPSE Ti2 microscope. Imaging parameters (e.g., laser power, gain) were defined for each primary antibody and were kept consistent between all sections in that primary antibody group.

### Image Analysis

Images for analysis of RNAscope sections were taken on a Keyence BZ-X800 microscope (KEYENCE; Osaka, Japan) at 40X magnification. Two images were acquired of each mouse LEC hemisphere, and 4 sections were imaged per mouse (total: ∼8 images/experiment for an n=3). The acquisition parameters were kept constant throughout imaging of all sections. Four fluorescent channels were used simultaneously; (1) the green channel was assigned for VIVID 520 dye (human APP probe), (2) the blue channel was used for DAPI nuclear stain, (3) the red channel was assigned for VIVID 570 dye (mouse APP probe), and (4) the far-red channel was assigned for VIVID 650 dye (CaMKIIα or Pvalb probes). A z-stack was taken (with 1 μm steps) of each hemisphere, and the full focus feature in the Keyence BZ-X800 analysis software was applied to compress each 10 μm z-stack. These compressed z-stacks were then used for image analysis in HALO v3.6 FISH-IF v2.1.4 (Indica Labs). For immunohistochemistry experiments, 2 sections were imaged per antibody per mouse (total: ∼4 images/experiment for an n=3-5). The acquisition parameters were kept constant throughout imaging of all sections. Two channels were used simultaneously; (1) the green channel was assigned for CaMKII.eYFP expression, (2) the red channel was assigned for Alexa Fluor^TM^ 647. For each slice, four line scans of 200 thickness were analyze from the pia to the end of the 60x photo (∼180-200 μm total) using ImageJ software. Antibody brightness was normalized to CaMKII.eYFP expression for each condition to control for any slight variation in viral expression. The obtained data was then analyzed for figure generation in Prism (GraphPad). The process from sample fixation to image analysis covered a four-day time frame.

### Flow Cytometry

Flow cytometry was performed for the confirmation of viral hAPP protein expression. For positive control 5xFAD and their wild-type littermates were used and injected stereotactically in the somatosensory cortex with 1:1 ratio of AAV(PHP.eB).E2.GFP and saline (0.3 µL). While for hAPP expression C57Bl/6J were also stereotactically injected at a 1:1 ratio with AAV(PHP.eB).E2.GFP and saline or AAV(PHP.eB).hAPP (0.3 µL total). 2-3 weeks after the injections, mice were euthanized (8–10 weeks old) by decapitation and acute slices of 250 µm were obtained and then micro-dissected to isolate the cortical region containing GFP^+^ expressing cells using an epi-fluorescent stereoscope (Olympus SZX12). Virus-injected microdisected regions of the brain were then placed into a cutting solution with 0.5 mg/mL protease (P5147– 100MG, Sigma-Aldrich) for 60 minutes with continuous carbogen gas bubbling. Samples were then manually triturated in 300 µL of 1% PBS into a single-cell suspension. Cells were stained with a human-specific APP antibody (SIG-39320, Biolegend). Cells were first fixed in 1x fixation buffer (eBioscience; cat# 00-8222-49) for 30 min on ice, then washed 3x in PBS. Cells were then permeabilized for 30 min using 1x permeabilization buffer (eBioscience; cat# 00-8333-56) on ice. To determine the presence of hAPP in these cells, fixed and permeabilized cells were incubated with human-specific APP antibody (SIG-39320, Biolegend) at dilution 1:250 for 1 h. cells were then washed with permeabilization buffer 3 times and incubated for 30 minutes with a secondary antibody at dilution 1:500. Cells were finally washed 3 times with permeabilization buffer, as mentioned above. After the last wash, 250 µl of PBS was added, vortexed and kept on ice in the dark until flow cytometry was performed. Compensation was performed prior to the experiment using a single fluorophore labelled OneComp eBeads™ Compensation Beads (Catalog number: 01-1111-41; Thermofisher).

### Statistics and Analysis

Custom python scripts, Axograph, Graphpad Prism (Graphpad Software), and Excel (Microsoft) were used for analysis with values in text and figures. Statistical differences were deemed significant with α values of p < 0.05. Two-tailed unpaired and paired t-tests were used for unmatched and matched parametric datasets, respectively. Where appropriate, group data were compared with 1 or 2-way ANOVA and significance between groups noted in figures was determined with Tukey’s or Sidak’s multiple post-hoc comparison tests. Normality was determined using D’Agostino & Pearson omnibus or Shapiro-Wilk tests. For permutation-based false discovery proportion (FDP) estimation, we used permFDP package in R [version 4.3.1], https://github.com/steven-shuken/permFDP for correcting p-values for multiple hypothesis testing. Using a family of unadjusted p values, and 500 permutations, this function returned the corrected rejection threshold to control false discovery (FDR<5%). For this dataset, the adjusted p-value threshold was 0.0431.

### K-means clustering and Principal Component Analysis

K-means clustering and Principal component analysis (PCA) were conducted on datasets from excitatory neurons and PV interneurons, respectively, in the LEC. All passive and active properties were used for each cell to conduct unsupervised clustering. Post-clustering and analysis, Ctrl or hAPP identities were restored to their respective cells.

### Analyses of MS data and bioinformatics analyses

Within each MS study, we compared bulk proteomes to SA-enriched proteomes to confirm that expected proteins (from either PV-INTs) were indeed enriched while nonneuronal proteins (e.g. glial proteins) were de-enriched as compared to bulk brain proteomes. We also identified proteins unique to bulk or SA-enriched pulldown samples. Within SA-enriched biotinylated proteins, we restricted our analyses to those proteins that were confidently biotinylated and enriched (based on statistical significance unadj. P<0.05 as well as 2-fold enrichment in biotinylated vs. non biotinylated samples). This allowed us to exclude proteins that were non-specifically enriched by streptavidin beads. Within biotinylated proteins, group comparisons were performed using a combination of approaches, including differential abundance analysis, hierarchical clustering analysis (Broad Institute, Morpheus, https://software.broadinstitute.org/morpheus), as well as PCA, (in SPSS Ver 26.0 or R). Differential abundance analyses were performed on log2 transformed and normalized abundance values using two-tailed unpaired T-test for 2 groups assuming equal variance across groups or one-way ANOVA + post-hoc Tukey HSD tests for >2 groups). Unadjusted and FDR-corrected comparisons were performed, although we relied on unadjusted p-values along with effect size (fold-enrichment) to improve stringency of analyses. After curating lists of differentially enriched proteins, gene set enrichment analyses (GSEA) were performed (AltAnalyze Ver 2.1.4.3) using all proteins identified across bulk and pulldown proteomes as the reference (background list). Protein-protein-interactions between proteins within lists of interest were examined using STRING (https://stringdb.org/cgi/input?sessionId=bqsnbjruDXP6&input_page_show_search=on<u>)123</u>. We also performed GSVA of DEPs identified in bulk as well as PV interneuron proteomes from SS Ctx and LEC to complement GSEA^85,86^. As previously published, statistical differences in enrichment scores for each ontology comparing two groups, were computed by comparing the true differences in means against a null distribution which was obtained by 1000 random permutations of gene labels. Benjamini & Hochberg false discovery rate adjusted p values <0.05 were considered significant. The reference gene sets for GSVA were the M5 (Mouse) Ontology Gene Sets from MSigDB (https://www.gseamsigdb.org/gsea/msigdb/mouse/collections.jsp?targetSpeciesDB=Mouse#M5).

### Data Availability

The PV-interneuron mass spectrometry proteomics data have been deposited to the ProteomeXchange Consortium via the PRIDE partner repository with the dataset identifier PXD053491. The 2020 mouse UniProt database (downloaded from https://www.uniprot.org/help/reference_proteome) was used for searches of mass spectrometry data. Source data are provided with this paper.

#### Project Name

Cortical region confers neuron-type-specific vulnerability to human APP expression

#### Project accession

PXD053491

## Acknowledgements

Contributions:

Conceptualization: AMG, MJR

Methodology and investigation: AMG, EB, PK, KEM, VJO, KS, CCR, AE, DD, NTS, SR, DW, MJR

Writing-Original draft: AMG, MJR

Writing-Review and Editing: MJR, AMG, SR, DW, CF

Funding acquisition: MJR, SR, AMG, EB

Resources: MJR, NTS, SR

---

**Extended Data Table 1.**
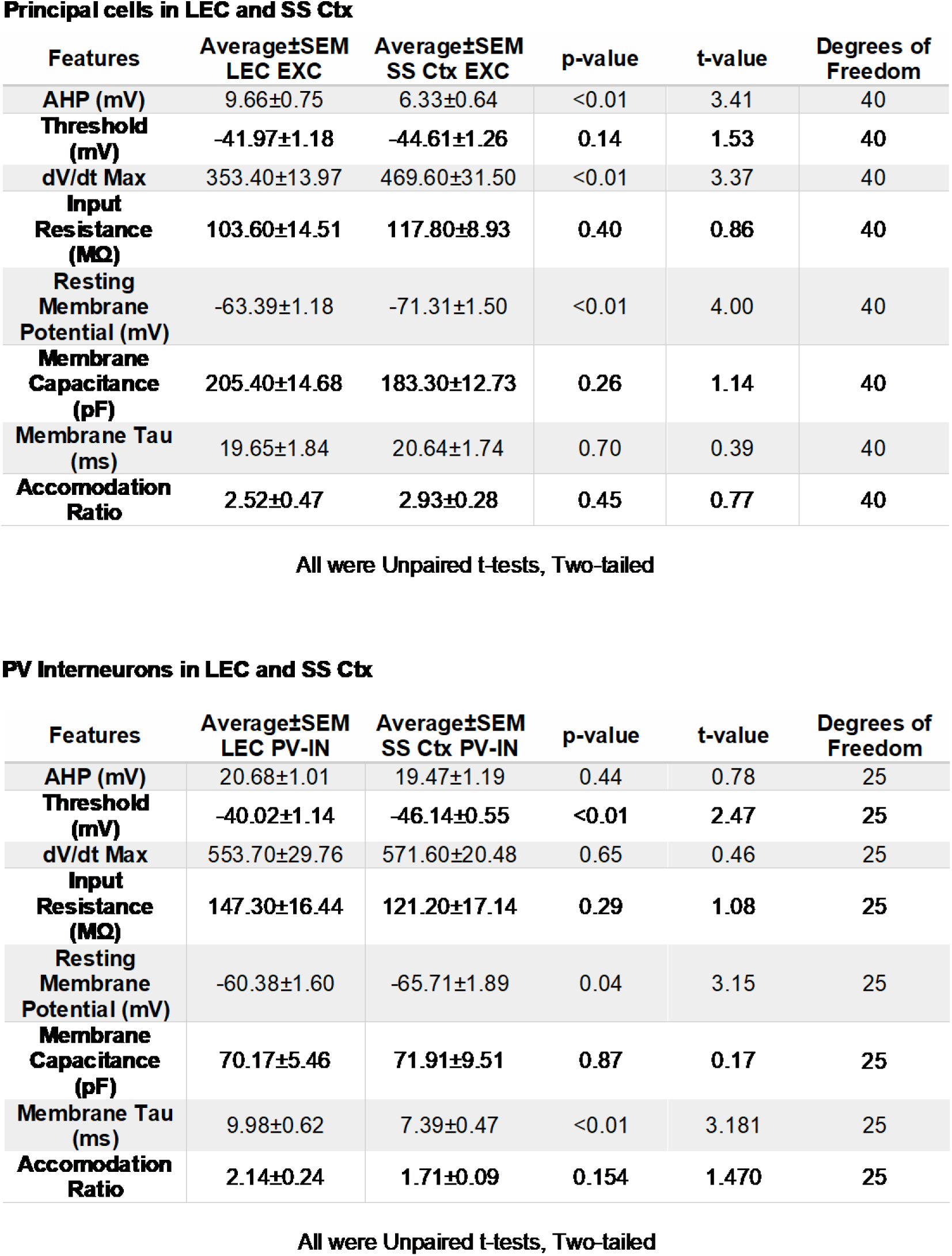
Passive and active properties of LEC and SS Ctx neurons. Passive and active properties of principal excitatory cells (top) or tdTom+ PV interneurons (bottom) in L2 LEC and L5 SS Ctx.

**Extended Data Figure 1.**
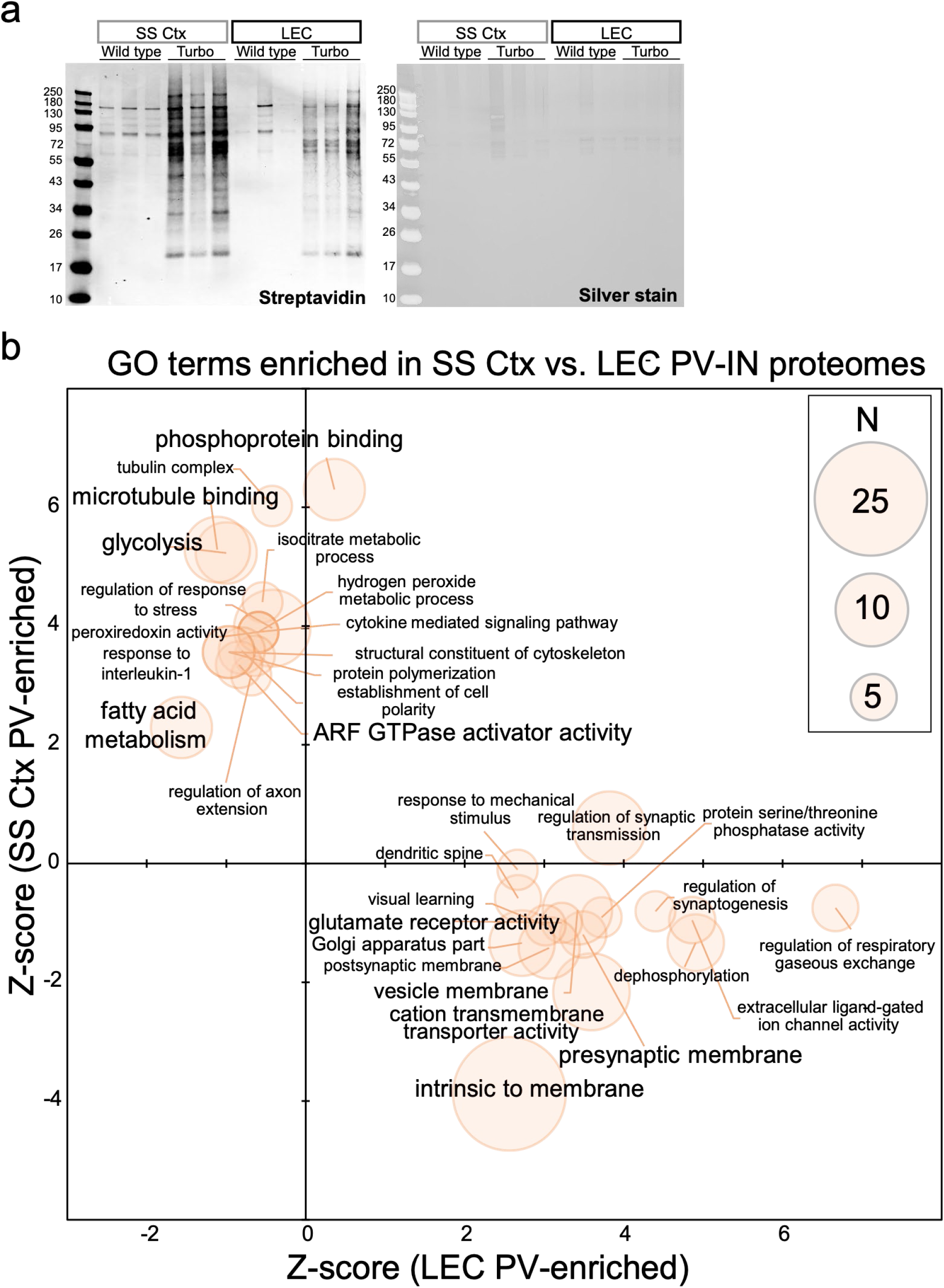
Enriched biotinylated PV-interneuron proteins from the SS Ctx and LEC are neuron-specific. **a.** Western blot (left) and silver stain (right) visualization of enriched biotinylated proteins in PV-interneurons (IN) from the SS Ctx and LEC after streptavidin-pulldown and elution of biotinylated proteins from a 10% aliquot of beads. **b.** GSEA of ³ 2-fold enriched biotinylated PV-interneuron proteins from the SS Ctx and LEC from PV-CIBOP mice, as compared to a reference protein list of both regions (n = 807) showed enrichment of synaptic and neuronal proteins confirming neuronal labeling. The orange dot size represents the number of gene symbols represented in each GO term. WT (Control) or Rosa26TurboID/wt (PV-CIBOP) mice (n=3 per genotype, including males and females).

**Extended Data Figure 2.**
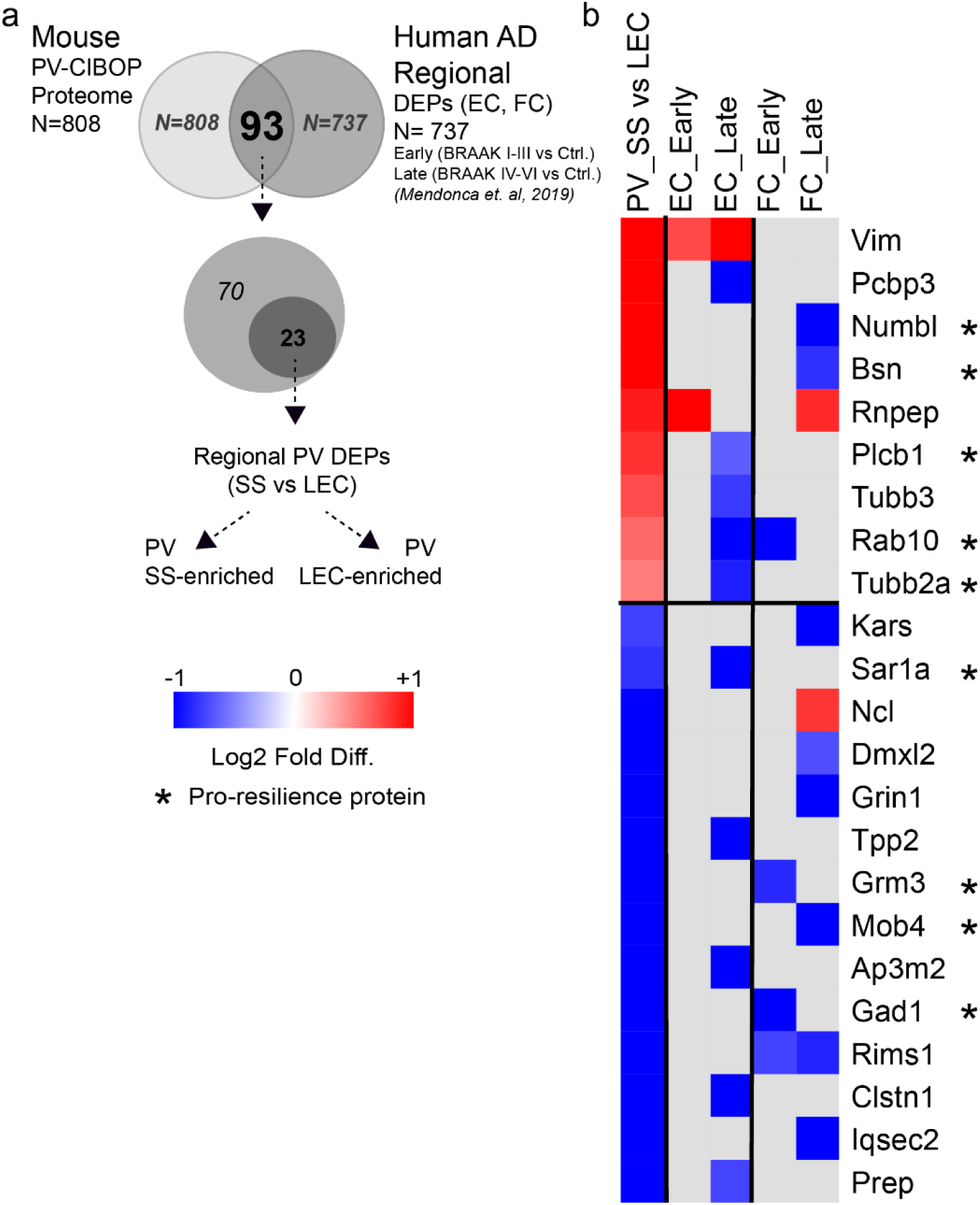
PV-CIBOP regional proteins that also demonstrate region-specific changes in human AD brain (related to Figure 1). **a.** Mouse PV-CIBOP regional proteins (SS Ctx vs. LEC) that were also identified as regional DEPs (AD vs. Ctrl) from human post-mortem brain regions (EC and FC)^23^. (EC: Entorhinal Cortex; FC: Frontal Cortex; Early: Braak I-III vs. Ctrl; Late: Braak IV-VI vs. Ctrl). See also Extended Data Datasheet 2 for related data and analyses.

**Extended Data Figure 3.**
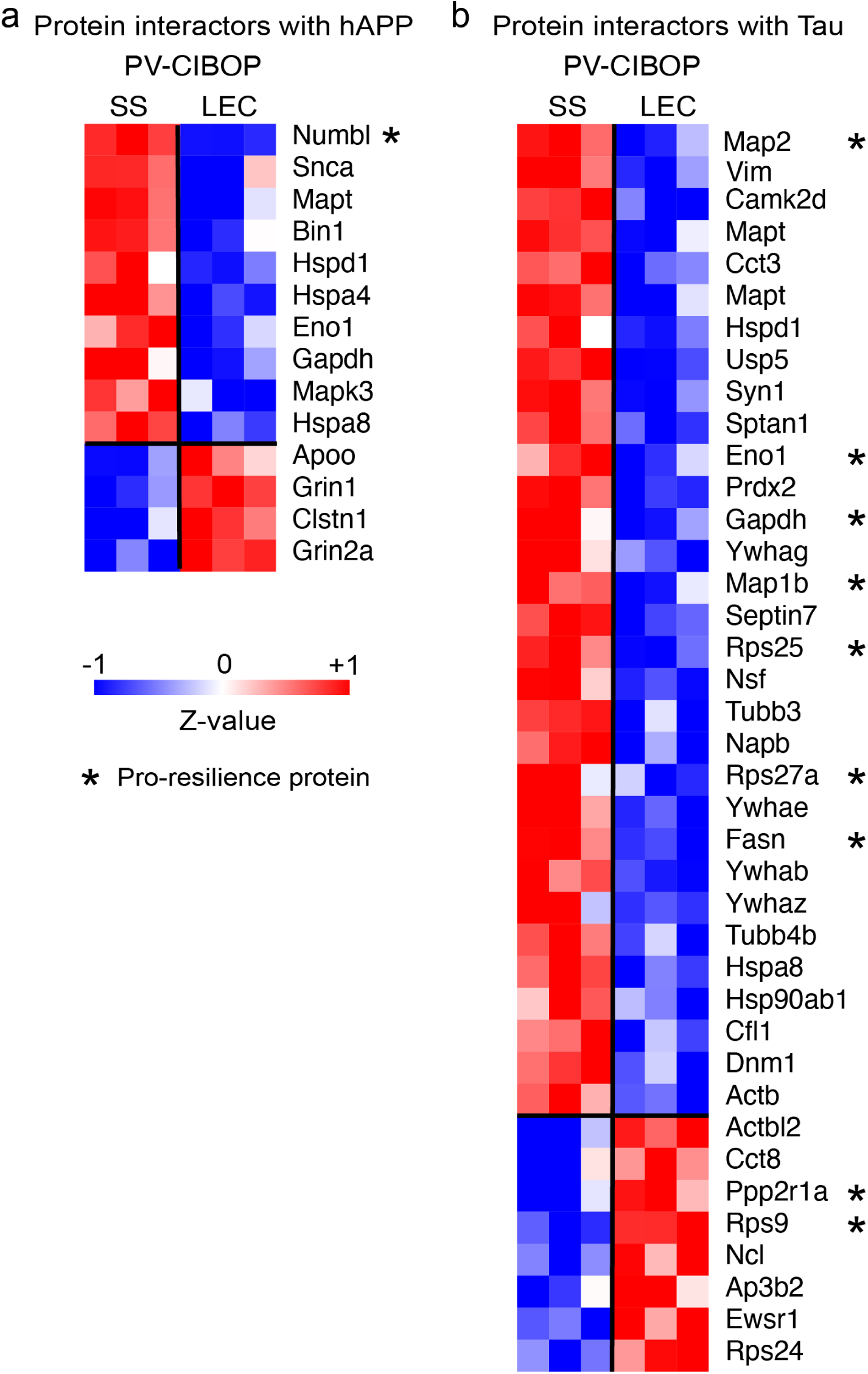
PV-CIBOP identified regional proteins with known interactions with human APP and Tau (Related to Figure 1). **a.** Heatmap representation of PV regional DEPs (SS Ctx vs. LEC) that are known protein-protein interactors with human APP (from 243 known APP interactors derived from the STRING database). Proteins marked with * indicate those that are also known pro-resilience proteins. As expected, the APP interactors were enriched in proteins involved in lipid binding, amyloid beta processing, cholesterol metabolism, and complement and coagulation cascade. **b.** Heatmap representation of PV interneuron regional DEPs (SS Ctx vs. LEC) that are known protein-protein interactors with Tau from human brain. Proteins marked with * indicate those that are also known pro-resilience proteins. Also see Extended Data Datasheets 3 and 4 for related results and analyses.

**Extended Data Figure 4.**
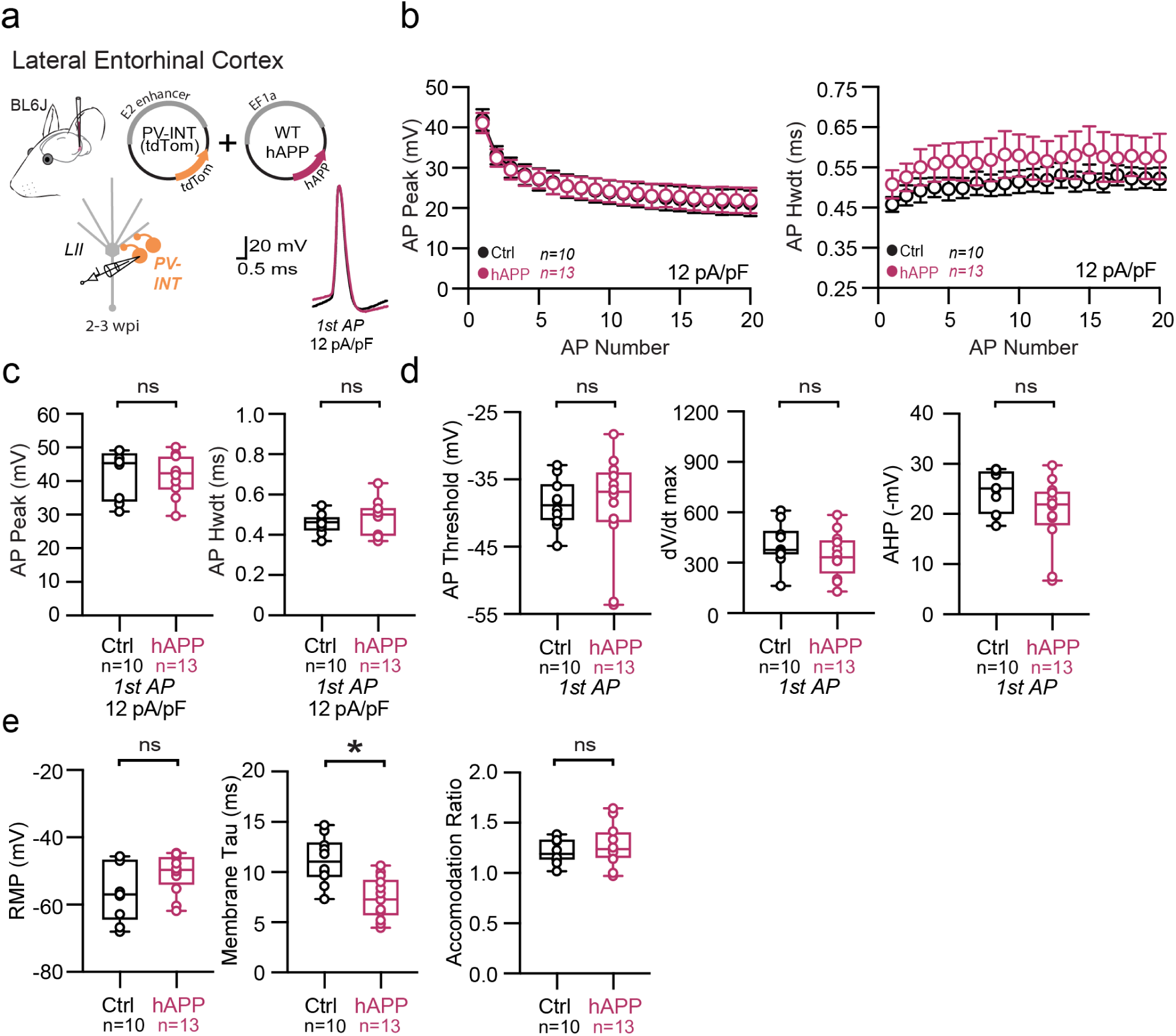
Passive and active properties of LEC PV interneurons after hAPP injection. **a.** Graphical summary of AAV.E2.tdTom and AAV.EF1a.hAPP (or for Ctrl, saline) stereotactic injection in the Lateral Entorhinal Cortex. tdTom+ PV interneurons were fluorescently targeted for whole-cell current-clamp recordings. AP waveforms of tdTom+ PV interneurons were compared at 12 pA/pF square pulse injections in WT mice from Ctrl and hAPP injected. Aps from the 1^st^ spike in the train are superimposed for comparison. **b.** Relationship between AP peak (p=0.88, df=21) or width (p<0.0001, df=21) in WT mice and AP # during spike trains elicited with a 12 pA/pF current injection. **c.** Summary data of AP properties. LEC PV interneurons after hAPP injection 1^st^ AP peak (p=0.94, t=0.08) and half-width remains unchanged (p=0.51, t=0.67, df=21). **d.** Summary data of AP properties. LEC PV interneurons after hAPP injection AP threshold (p=0.85, t=0.20), dV/dt max (p=0.12, t=1.6), and AHP (0.63, t=0.49) (df=21) remained unchanged. **e.** Summary data of AP properties. LEC PV interneurons after hAPP injection show unchanged Resting Membrane Potential (p=0.09, t=1.79) and Accomodation Ratio (p=0.66, t=0.44), but a reduction in Membrane Tau(p=0.0008, t=3.93) (df=21). For all summary graphs, data are expressed as mean (± SEM). For **b**: Statistical significance is denoted as *=p<0.05, as determined by Two-way ANOVA with Sidak’s multiple comparison test. For c, d, e: Individual data points and box plots are displayed. Statistical significance is denoted as *=p<0.05, as determined by two-tailed unpaired t-test.

**Extended Data Figure 5.**
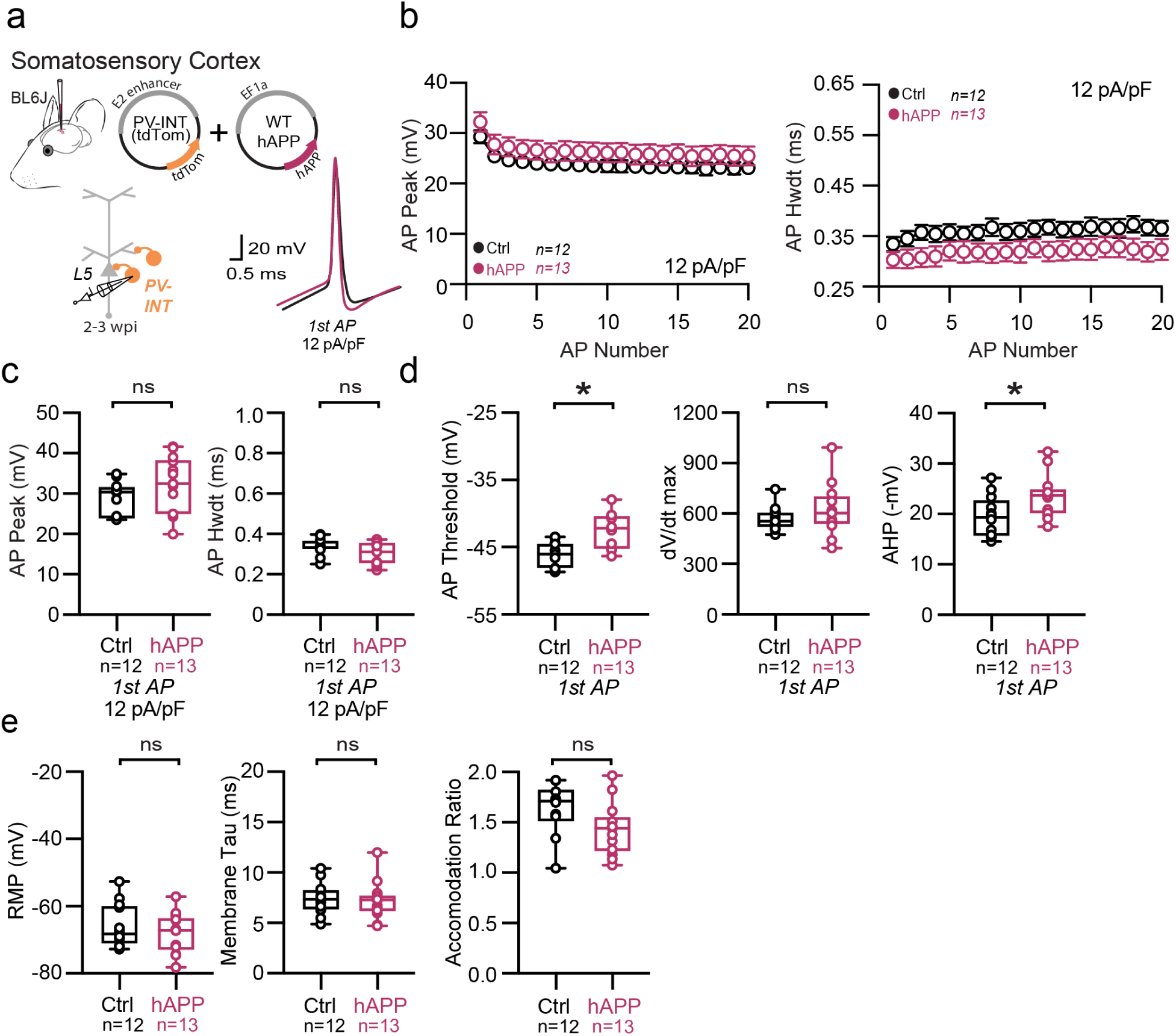
Passive and active properties of SS Ctx interneurons after hAPP injection. **a.** Graphical summary of AAV.E2.tdTom and AAV.EF1a.hAPP (or for Ctrl, saline) stereotactic injection in the SS Cortex. tdTom+ PV interneurons were fluorescently targeted for whole-cell current-clamp recordings. AP waveforms of tdTom+ PV interneurons were compared at 12 pA/pF square pulse injections in WT mice from Ctrl and hAPP injected. Aps from the 1^st^ spike in the train are superimposed for comparison. **b.** Relationship between AP peak (p=0.17, t=1.42 df=23) or width (p=0.15, t=1.49 df=23) for in WT mice and AP # during spike trains elicited with a 12 pA/pF current injection. **c.** Summary data of AP properties. SS PV interneurons after hAPP injection 1^st^ AP peak (p=0.17, t=1.42) and half-width (p=0.15, t=1.49) (df=23) remain unchanged. **d.** Summary data of AP properties. SS PV interneurons after hAPP injection dV/dt max remains unchanged (p=0.32, t=1.01). AP Threshold (p=0.001, t=3.84) and AHP (p=0.02, t=2.42)(df=23) significantly increase after hAPP injection. **e.** Summary data of AP properties. SS PV interneurons after hAPP injection show unchanged Resting Membrane Potential (p=0.40, t=0.85), Accomodation Ratio (p=0.08, t=1.84), and Membrane Tau (p=0.81, t=0.24) (df=23). For all summary graphs, data are expressed as mean (± SEM). For **b**: Statistical significance is denoted as *=p<0.05, as determined by Two-way ANOVA with Sidak’s multiple comparison test. For c, d, e: Individual data points and box plots are displayed. Statistical significance is denoted as *=p<0.05, as determined by two-tailed unpaired t-test.

**Extended Data Figure 6.**
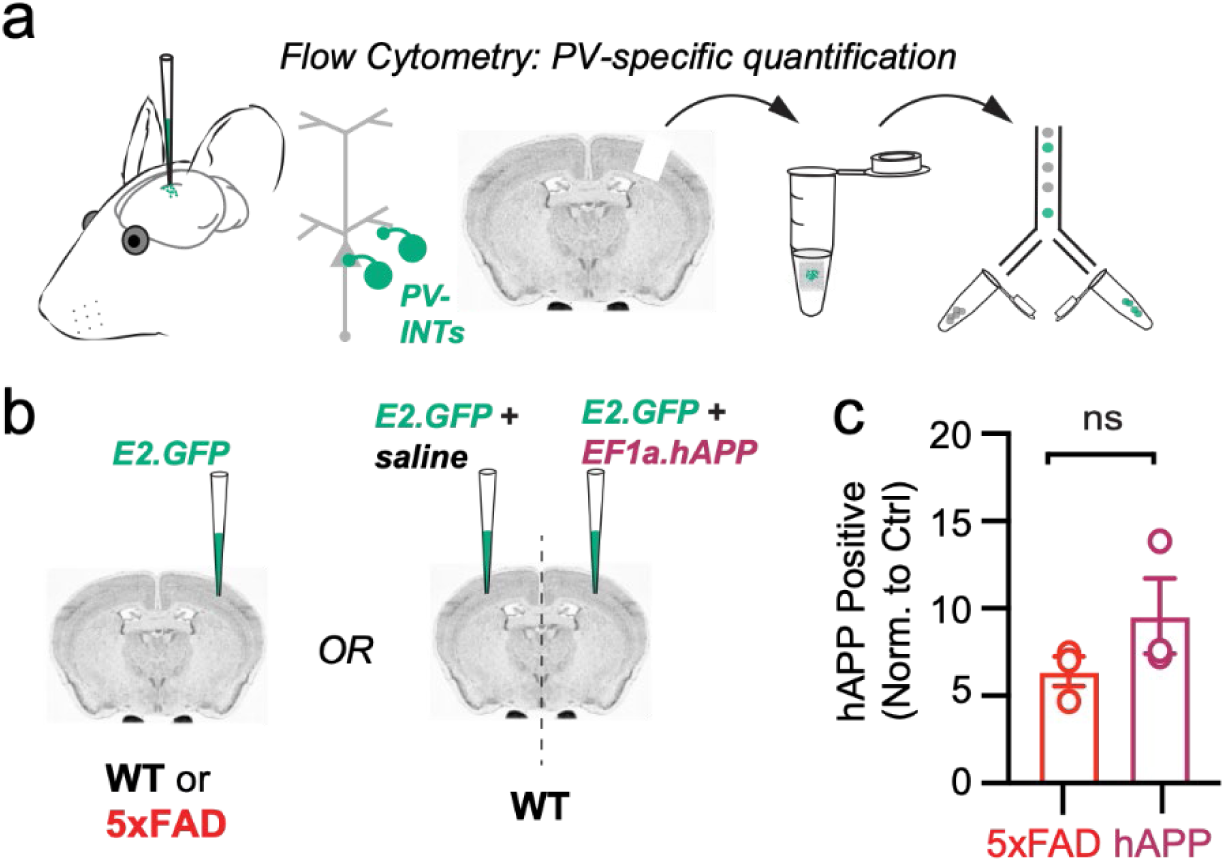
Confirmation of hAPP protein in PV interneurons using PV-specific flow cytometry. **a.** Graphical summary of PV-specific flow cytometry workflow. Region containing fluorescent PV interneurons after AAV.E2.GFP stereotactic injection in the SS Cortex was microdissected, triturated, and sorted based on GFP+ signal. Subsequent confirmation specifically for human APP was completed. **b.** WT and 5xFAD mice (∼ 2 months) were injected with AAV.E2.GFP and sorted using flow cytometry. WT mice were also used to compare AAV.E2.GFP + EF1a.hAPP injected SS Ctx to the contralateral hemisphere where EF1a was replaced with an equal volume of saline. **c.** Both groups were normalized to their control groups (WT littermates for 5xFAD; contralateral hemi for the hAPP control). The number hAPP expressing PV-interneurons did not significantly differ between 5xFAD and hAPP injected (p=0.24, t=1.37, df=4).

**Extended Data Figure 7.**
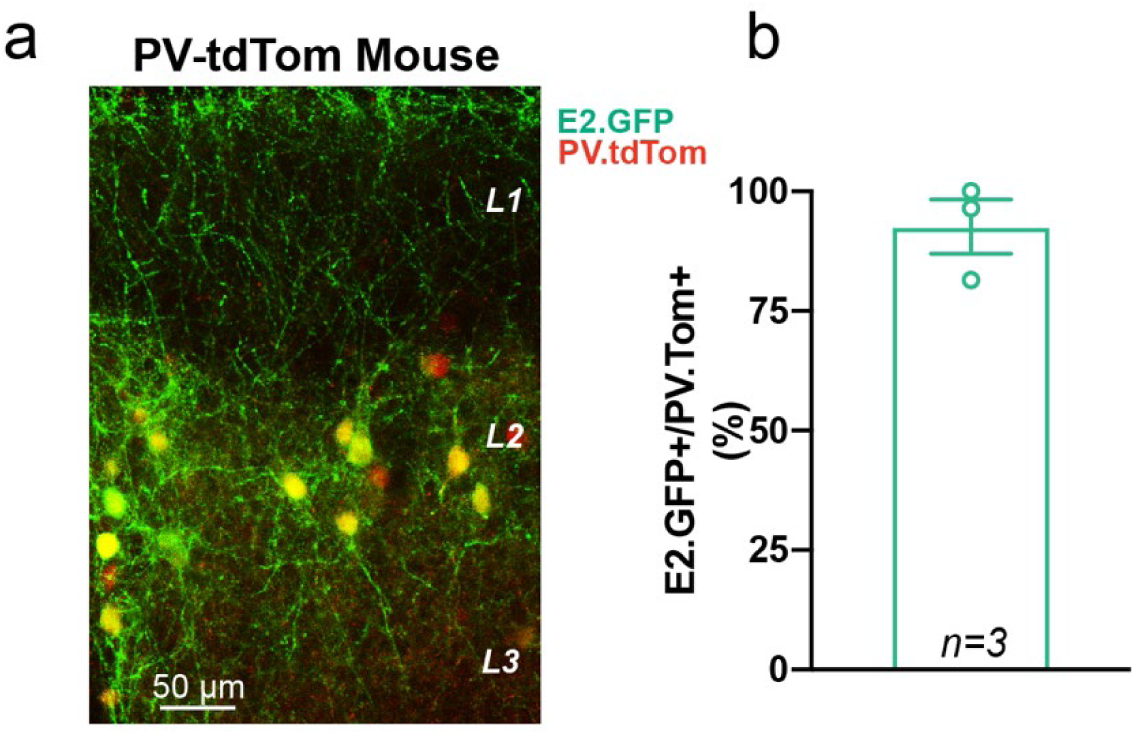
Specificity of AAV.pHP-eB.E2.GFP in the Lateral Entorhinal Cortex for stereotactic and retro-orbital injections. **a.** Stereotactic injection of AAV.pHP-eB.E2.GFP into the LEC of a PV-tdTom transgenic mouse. **b.** For three animals, three slices were analyzed for E2.GFP+ cells which were also PV.tdTom+ (Mean: 92.62 ± 5.7% for three biological replicates).

**Extended Data Figure 8.**
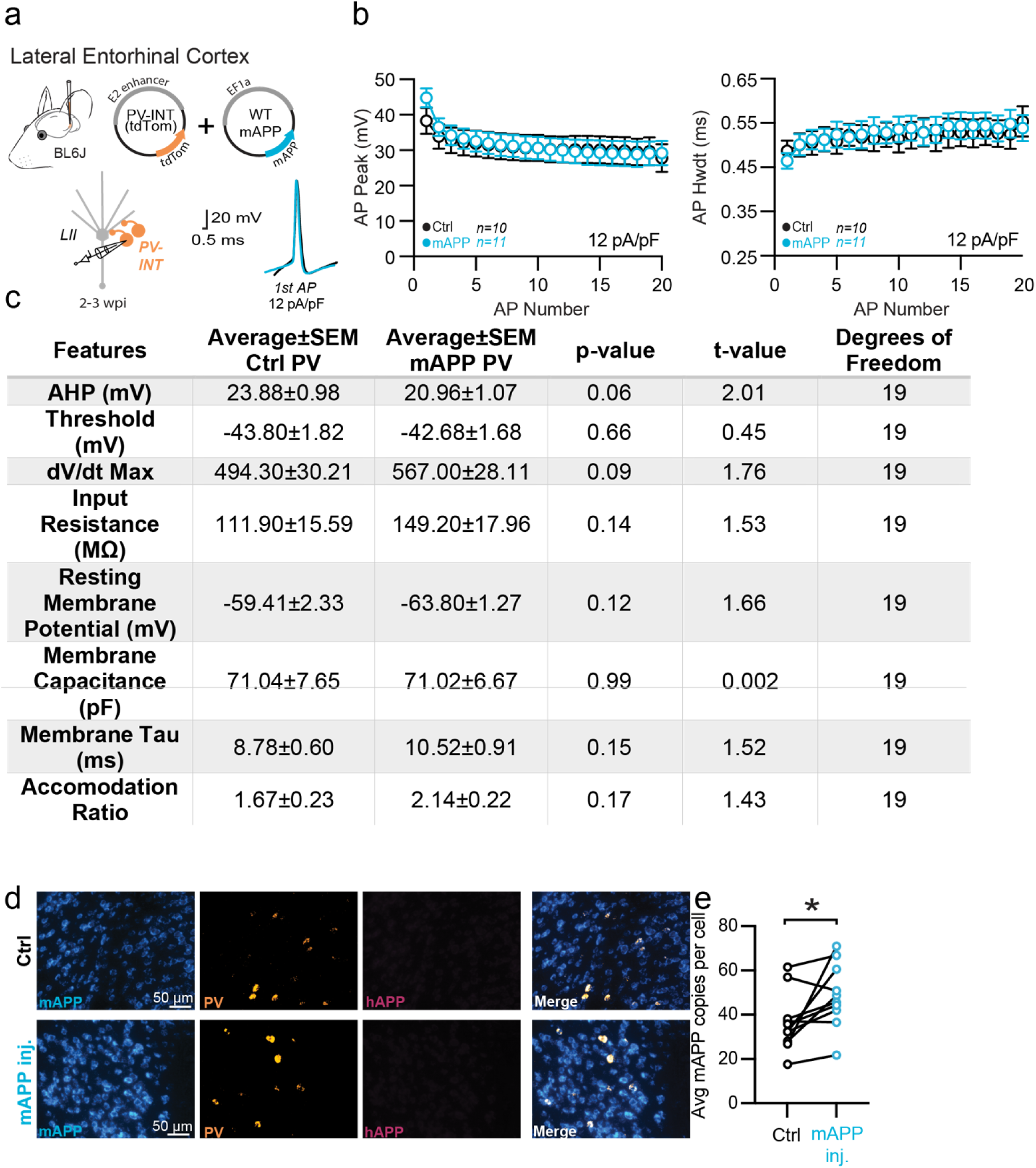
Passive and active properties of LEC PV interneurons after mAPP injection. **a.** Graphical summary of AAV.E2.tdTom and AAV.EF1a.mAPP (or for Ctrl, saline) stereotactic injection in the SS Cortex. tdTom+ PV interneurons were fluorescently targeted for whole-cell current-clamp recordings. AP waveforms of tdTom+ PV interneurons were compared at 12 pA/pF square pulse injections in WT mice from Ctrl and mAPP injected. Aps from the 1^st^ spike in the train are superimposed for comparison. **b.** Relationship between AP peak (p=0.61) or width (p=0.35) in WT mice and AP # during spike trains elicited with a 12 pA/pF current injection. **c.** Summary table of AP properties. **d.** RNAscope representative images at 40x magnification for Ctrl (top) and mAPP (bottom) injected mice: mAPP mRNA (cyan), Parvalbumin mRNA (gold), human APP mRNA (magenta), and a final merged image. **e.** RNAscope quantification for mAPP copies per DAPI+ cell comparing mAPP injected to the contralateral hemisphere endogenous mAPP expression. mAPP injected show a significant increase in mAPP copies in the injected hemisphere (p=0.03, t=2.57, df=9; two-tailed paired t-test). For all summary graphs, data are expressed as mean (± SEM). For **b**: Statistical significance is denoted as *=p<0.05, as determined by Two-way ANOVA with Sidak’s multiple comparison test. For c, e: statistical significance is denoted as *=p<0.05, as determined by two-tailed unpaired t-test.

**Extended Data Figure 9.**
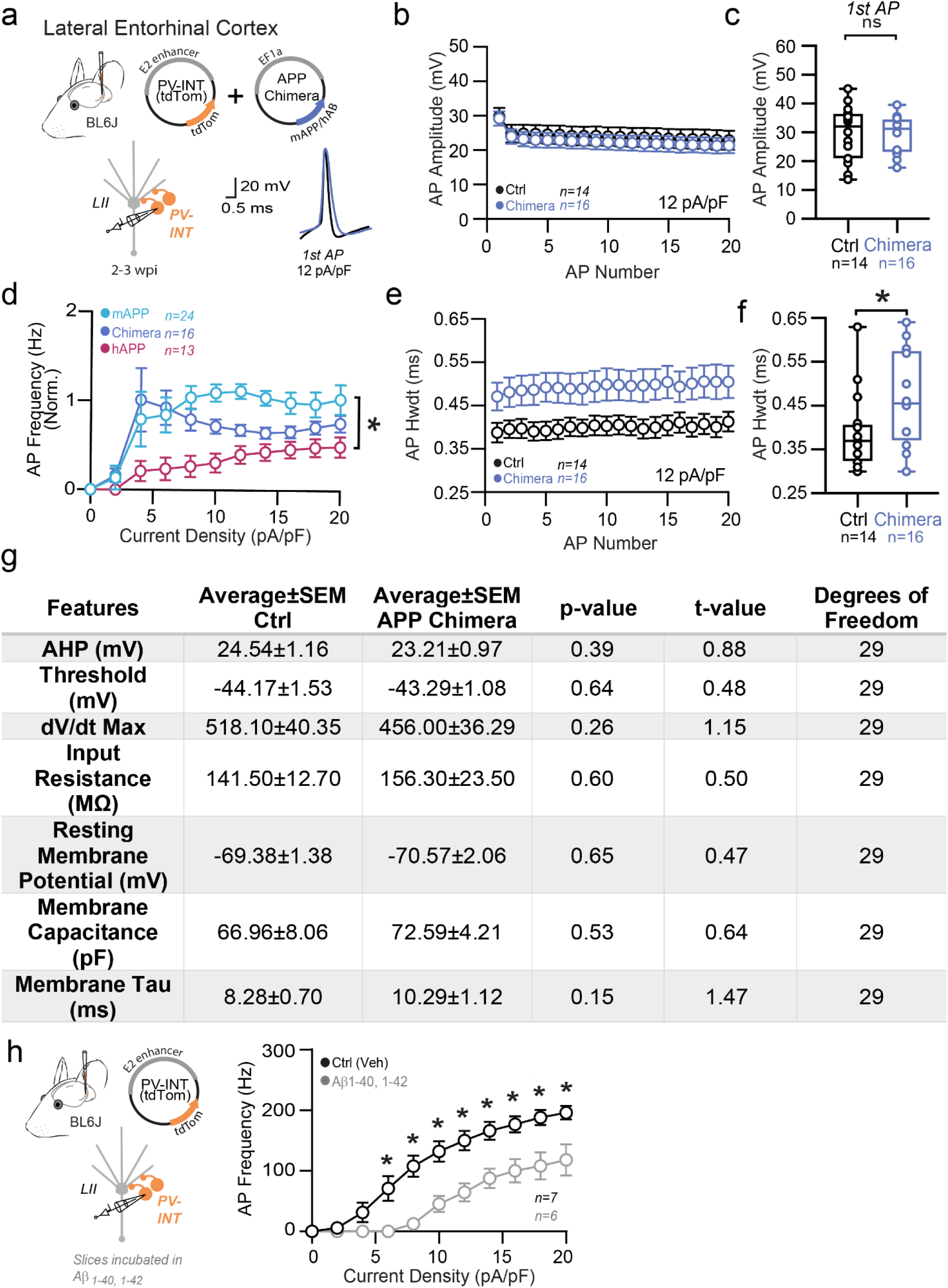
Passive and active properties of LEC PV interneurons after mAPP/hAB Chimera injection. **a.** Graphical summary of AAV.E2.tdTom and AAV.EF1a.mAPP/hAB Chimera (or for Ctrl, saline) stereotactic injection in the SS Cortex. tdTom+ PV interneurons were fluorescently targeted for whole-cell current-clamp recordings. AP waveforms of tdTom+ PV interneurons were compared at 12 pA/pF square pulse injections in WT mice from Ctrl and mAPP injected. Aps from the 1^st^ spike in the train are superimposed for comparison. **b.** Relationship between AP peak (b) or width (e) in WT mice and AP # during spike trains elicited with a 12 pA/pF current injection. **c.** Summary data of AP properties. LEC PV cells after Chimera injection 1^st^ AP peak (29.74 ± 2.61 mV, 29.18 ± 2.08 mV, hAPP and Ctrl respectively, p=0.87, t=0.17, df=24) remains unchanged. **d.** Group data summary of AP firing frequency in L2 LEC from mAPP injected (blue), Chimera injected (purple), and hAPP injected mice (magenta), all normalized to their dataset controls. **e.** Relationship between AP width in WT mice and AP # during spike trains elicited with a 12 pA/pF current injection. **f.** AP half-width displays a significant increase in the chimera group (0.39± 0.02 ms, 4.80 ± 0.03 ms, hAPP and Ctrl respectively, p=0.04, t=2.14, df=24). **g.** Summary table of effects of the mAPP/hAβ chimera. **h.** Summary of AP firing frequency in L2 PV interneurons from LEC Aβ incubated slices and vehicle controls. For all summary graphs, data are expressed as mean (± SEM). For c, f, g: statistical significance is denoted as *=p<0.05, as determined by two-tailed unpaired t-test.

**Extended Data Figure 10.**
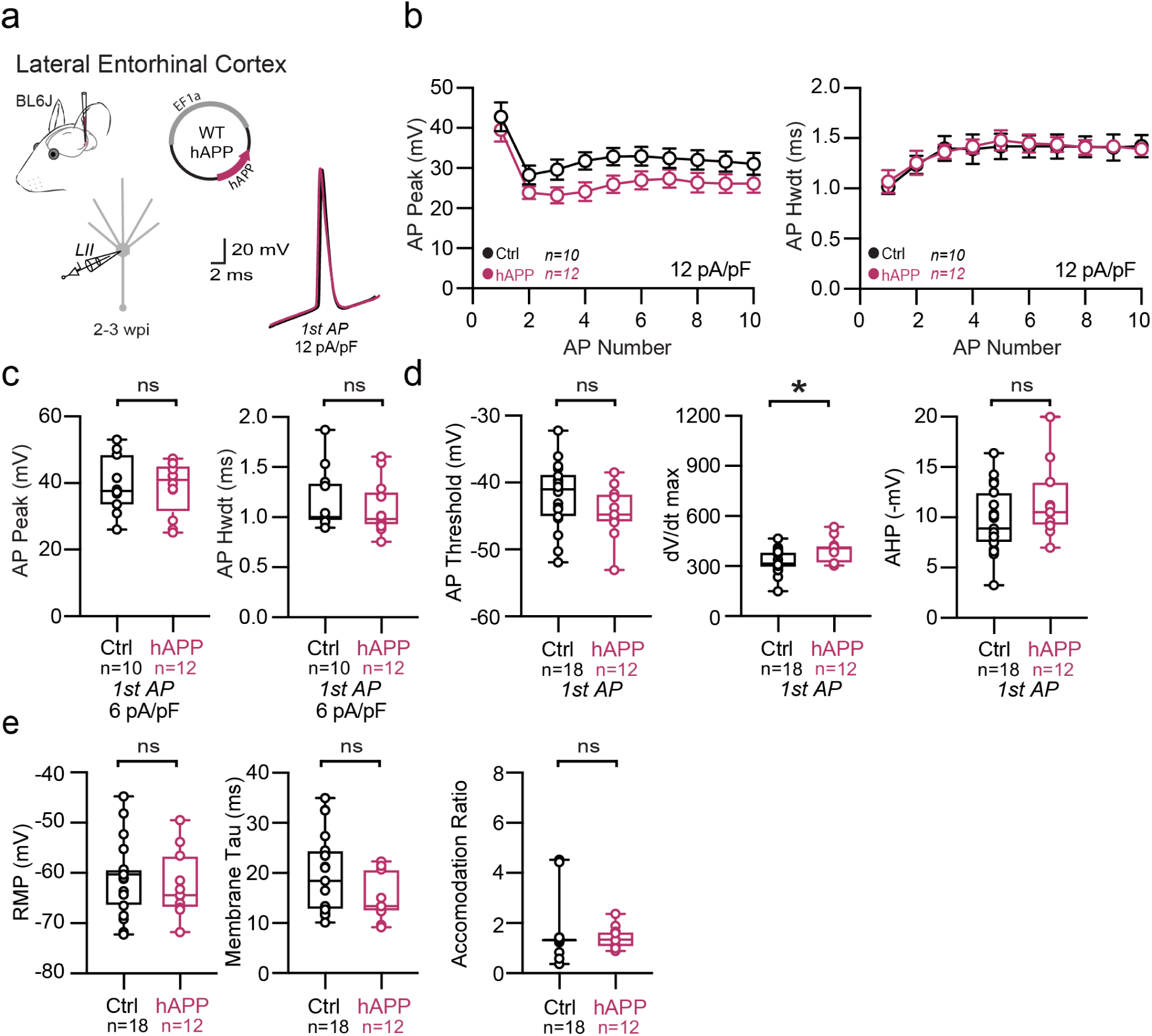
Passive and active properties of LEC excitatory neurons after hAPP injection. **a.** Graphical summary of AAV.EF1a.hAPP (or for Ctrl, saline) stereotactic injection in the LEC. Excitatory cells were targeted for whole-cell current-clamp recordings. AP waveforms of excitatory cells were compared at 12 pA/pF square pulse injections in WT mice from Ctrl and hAPP injected. Aps from the 1^st^ spike in the train are superimposed for comparison. **b.** Relationship between AP peak or width in WT mice and AP # during spike trains elicited with a 12 pA/pF current injection. **c.** Summary data of AP properties. LEC excitatory cells after hAPP injection 1^st^ AP peak (29.36 ± 4.14 mV, 28.62 ± 2.34 mV, hAPP and Ctrl respectively, p=0.87, t=0.17, df=28) and half-width remains unchanged (1.26 ± 0.10 ms, 1.50 ± 0.13 ms, hAPP and Ctrl respectively, p=0.23, t=1.28, df=28). **d.** Summary data of AP properties. LEC excitatory cells after hAPP injection AP threshold (p=0.18, t= 1.38, df = 28) and AHP remain unchanged (p=0.16, t=1.46, df=28), but dV/dt max shows a significance increase (399.7 ± 21.82, 331.0 ± 16.71), hAPP and Ctrl respectively, p=0.02, t=2.49, df=28). **e.** Summary data of AP properties. LEC excitatory cells after hAPP injection show unchanged Resting Membrane Potential (p=0.60, t=0.53, df=28), Accomodation Ratio (p=0.69, t=0.40, df=28), and Membrane Tau (p=0.08, t=1.84, df=28). For all summary graphs, data are expressed as mean (± SEM). For **b**: Statistical significance is denoted as *=p<0.05, as determined by Two-way ANOVA with Sidak’s multiple comparison test. For c, d, e: Individual data points and box plots are displayed. Statistical significance is denoted as *=p<0.05, as determined by two-tailed unpaired t-test.

**Extended Data Figure 11.**
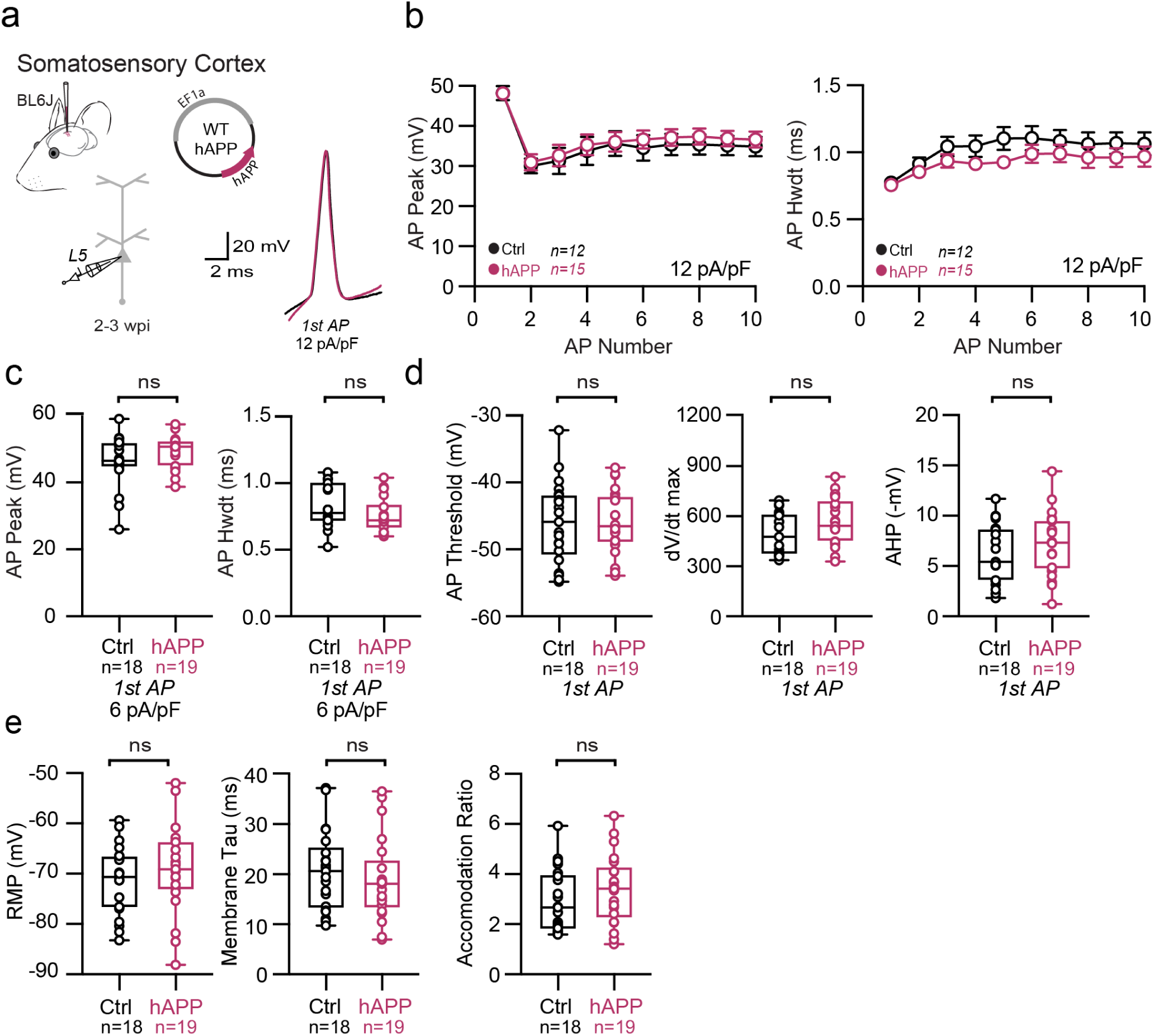
Passive and active properties of SS Ctx excitatory neurons after hAPP injection. **a.** Graphical summary of AAV.EF1a.hAPP (or for Ctrl, saline) stereotactic injection in the SS Ctx. Excitatory cells were targeted for whole-cell current-clamp recordings. AP waveforms of excitatory cells were compared at 12 pA/pF square pulse injections in WT mice from Ctrl and hAPP injected. Aps from the 1^st^ spike in the train are superimposed for comparison. **b.** Relationship between AP peak or width in WT mice and AP # during spike trains elicited with a 12 pA/pF current injection. **c.** Summary data of AP properties. SS Ctx excitatory cells after hAPP injection 1^st^ AP peak (p=0.19, t=1.34) and half-width (p=0.17, t=1.41) (df=40) remains unchanged. **d.** Summary data of AP properties. SS Ctx excitatory cells after hAPP injection AP threshold (p=0.97, t=0.04, df=40), dV/dt max (p=0.08, t=1.79, df=40), and AHP remain unchanged (p=0.32, t=1.00, df=40). **e.** Summary data of AP properties. SS Ctx excitatory cells after hAPP injection show unchanged Resting Membrane Potential (p=0.37, t=0.90, df=40), Accomodation Ratio (p=0.25, t=1.16, df=40), and Membrane Tau(p=0.48, t=0.71, df=40). For all summary graphs, data are expressed as mean (± SEM). For **b**: Statistical significance is denoted as *=p<0.05, as determined by Two-way ANOVA with Sidak’s multiple comparison test. For c, d, e: Individual data points and box plots are displayed. Statistical significance is denoted as *=p<0.05, as determined by two-tailed unpaired t-test.

**Extended Data Figure 12.**
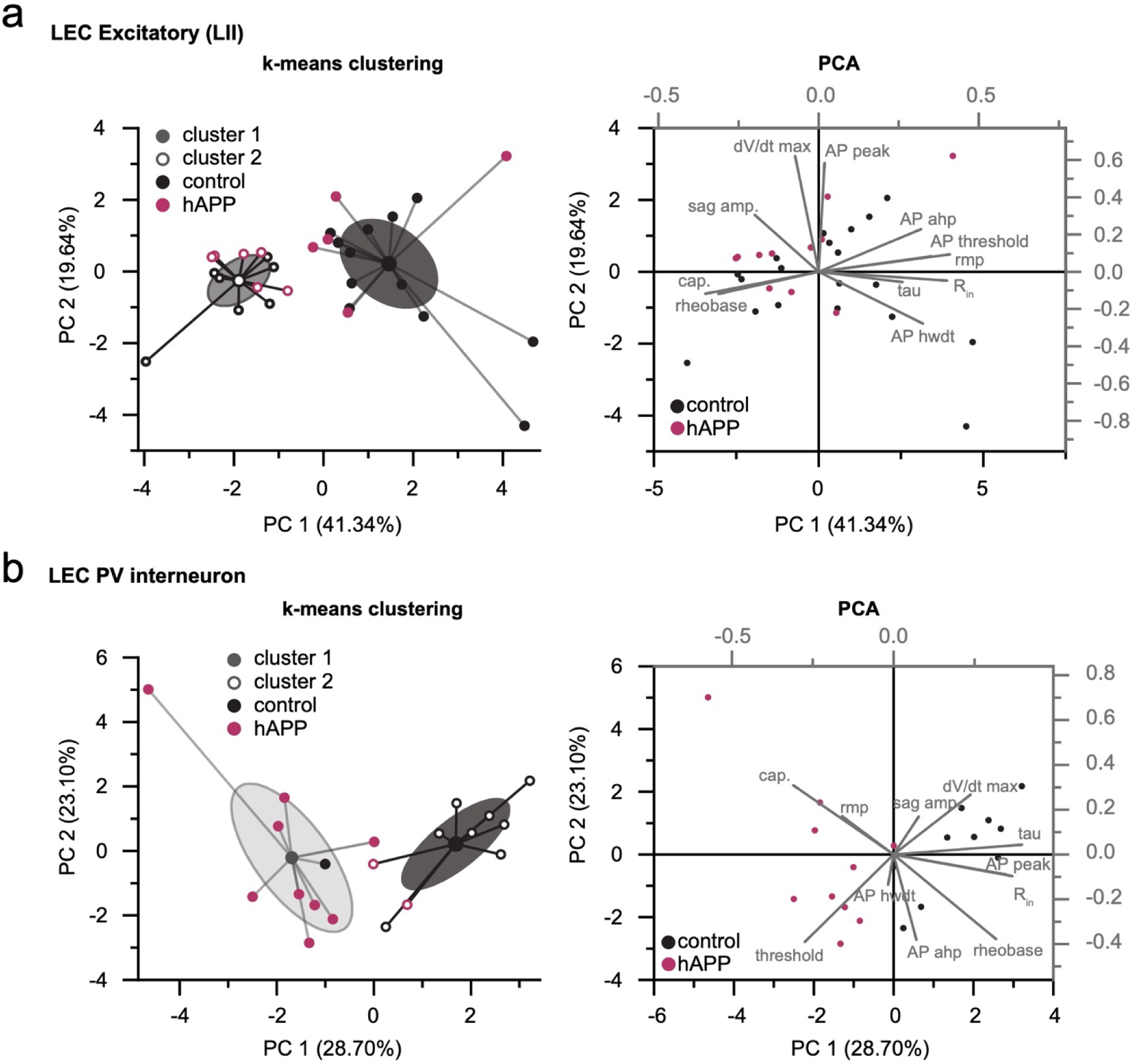
PCA analysis of LEC cell populations after hAPP injection. K-means clustering and Principal component analysis (PCA) plot performed on all cells based on active and passive properties for (a) excitatory cells and (b) PV interneurons. a. K-mean clustering fails to separate LEC excitatory cells based on hAPP identity (left), due to largely homogeneous active and passive properties, suggesting differences along different axes. b. Unsupervised clustering preserves hAPP identity in LEC PV interneurons.

**Extended Data Figure 13.**
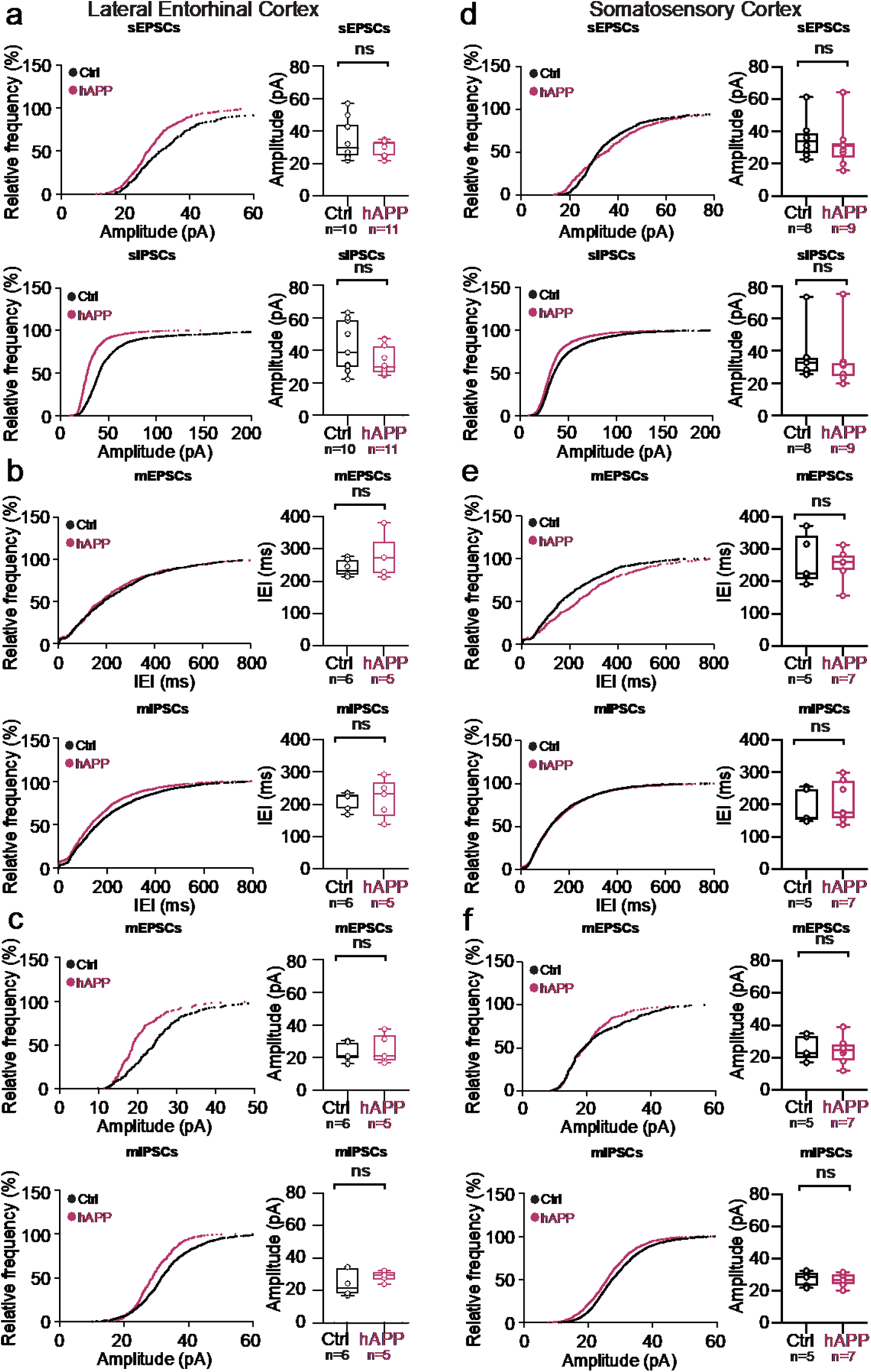
Adult-onset human APP does not alter mini frequencies or event kinetics in the LEC or SS Ctx. **a. Top:** Cumulative distribution curve for spontaneous EPSCs in the LEC showing the relationship of relative frequency of events to the amplitude (left). Quantified averages of event amplitude are displayed for each cell as individual data points and compared between Ctrl (black) and hAPP injected (magenta) conditions (right). L2 LEC sEPSCs showed no amplitude change (p=0.34, t=0.97, df=19). **Bottom:** Cumulative distribution curve for spontaneous IPSCs in the LEC showing the relationship of relative frequency of events to the amplitude (left). Quantified averages of event amplitude are displayed for each cell as individual data points and compared between Ctrl (black) and hAPP injected (magenta) conditions (right). L2 LEC sIPSCs show an unchanged amplitude (p=0.11, t=1.66, df=19). **b. Top:** Cumulative distribution curve for miniature EPSCs in the LEC showing the relationship of relative frequency of events to the inter-event interval (left). Quantified averages of IEIs are displayed for each cell as individual data points and compared between Ctrl (black) and hAPP injected (magenta) conditions (right). L2 LEC mEPSCs show no change in the IEIs (p=0.28, t=1.16, df=9). **Bottom:** Cumulative distribution curve for miniature IPSCs in the LEC showing the relationship of relative frequency of events to the IEIs (left). Quantified averages of IEIs are displayed for each cell as individual data points and compared between Ctrl (black) and hAPP injected (magenta) conditions (right). L2 LEC mIPSCs show no change in the IEIs (p=0.80, t=0.27, df=9). **c. Top:** Cumulative distribution curve for miniature EPSCs in the LEC showing the relationship of relative frequency of events to the amplitude (left). Quantified averages of event amplitude are displayed for each cell as individual data points and compared between Ctrl (black) and hAPP injected (magenta) conditions (right). L2 LEC mEPSCs no amplitude change (p=0.67, 0.45, df=9). **Bottom:** Cumulative distribution curve for miniature IPSCs in the LEC showing the relationship of relative frequency of events to the amplitude (left). Quantified averages of event amplitude are displayed for each cell as individual data points and compared between Ctrl (black) and hAPP injected (magenta) conditions (right). L2 LEC mIPSCs show an unchanged amplitude (p=0.27, t=1.18, df=9). **d. Top:** Cumulative distribution curve for spontaneous EPSCs in the SS Ctx showing the relationship of relative frequency of events to the amplitude (left). Quantified averages of event amplitude are displayed for each cell as individual data points and compared between Ctrl (black) and hAPP injected (magenta) conditions (right). L5 SS Ctx sEPSCs no amplitude change (p= 0.57, t=0.59, df=15). **Bottom:** Cumulative distribution curve for spontaneous IPSCs in the SS Ctx showing the relationship of relative frequency of events to the amplitude (left). Quantified averages of event amplitude are displayed for each cell as individual data points and compared between Ctrl (black) and hAPP injected (magenta) conditions (right). L5 SS Ctx sIPSCs show an unchanged amplitude (p=0.75, t=0.33, df=15). **e. Top:** Cumulative distribution curve for miniature EPSCs in the SS Ctx showing the relationship of relative frequency of events to the IEIs (left). Quantified averages of IEIs are displayed for each cell as individual data points and compared between Ctrl (black) and hAPP injected (magenta) conditions (right). L2 LEC mEPSCs show no change in the IEIs (p=0.74, t=0.35, df=10). **Bottom:** Cumulative distribution curve for miniature IPSCs in the SS Ctx showing the relationship of relative frequency of events to the IEIs (left). Quantified averages of IEIs are displayed for each cell as individual data points and compared between Ctrl (black) and hAPP injected (magenta) conditions (right). L5 SS Ctx mIPSCs show no change in the IEIs (p=0.65, t=0.47, df=10; mEPSC: p=0.74, t=0.35, df=10). **f. Top:** Cumulative distribution curve for miniature EPSCs in the SS Ctx showing the relationship of relative frequency of events to the amplitude (left). Quantified averages of event amplitude are displayed for each cell as individual data points and compared between Ctrl (black) and hAPP injected (magenta) conditions (right). L5 SS Ctx mEPSCs no amplitude change (p=0.77, t=0.31, df=10). **Bottom:** Cumulative distribution curve for miniature IPSCs in the SS Ctx showing the relationship of relative frequency of events to the amplitude (left). Quantified averages of event amplitude are displayed for each cell as individual data points and compared between Ctrl (black) and hAPP injected (magenta) conditions (right). L5 SS Ctx mIPSCs show an unchanged amplitude (p=0.78, t=0.29, df=10). For a, b, c, d, e, f: Individual data points and box plots are displayed. Statistical significance is denoted as *=p<0.05, as determined by two-tailed unpaired t-test

**Extended Data Figure 14.**
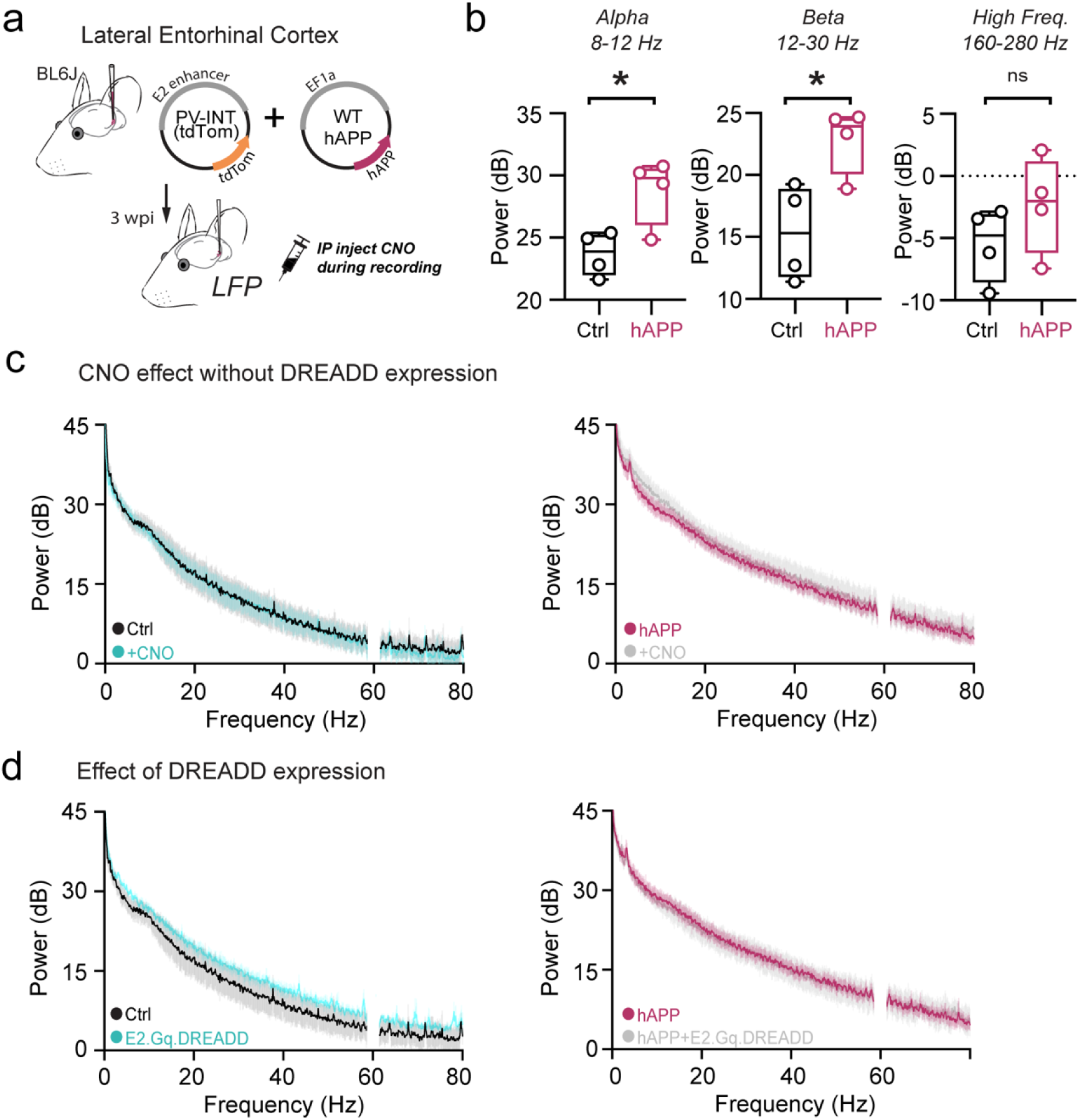
DREADD expression or CNO injections alone do not impact LEC circuit excitability. **a.** Graphical summary of AAV.EF1a.hAPP (or for Ctrl, E2.tdTom) stereotactic injection in the Lateral Entorhinal Cortex. **b.** Power [derived from PSD analysis] averaged for each mouse in the Alpha (8-12 Hz), Beta (12-30 Hz), High Frequency (160-280 Hz) frequency ranges and compared between Ctrl and hAPP. Alpha (Ctrl: 23.70± 0.89, Min. 21.64, Max. 25.40, Range. 3.76; hAPP: 28.79 ± 1.35, Min. 24.84, Max. 30.74, Range. 5.90; p=0.020, t=3.15, df=6); Beta (Ctrl: 15.32± 1.93, Min. 11.38, Max. 19.26, Range. 7.88; hAPP: 22.86 ± 1.35, Min. 18.90, Max. 24.67, Range. 5.77; p=0.019, t=3.20, df=6), High Frequency (Ctrl: −5.47± 1.50, Min. −9.43, Max. −2.87, Range. 6.56; hAPP: −2.34 ± 1.97, Min. −7.43, Max. 2.08, Range. 9.51; p=0.2530, t=1.26, df=6). Individual data points and box plots are displayed. Statistical significance is denoted as *=p<0.05, as determined by two-tailed unpaired t-test. **c.** Power Spectral Density [PSD] analysis to assess if CNO administration impacts excitability in the absence of DREADD expression (dark lines: mean, light lines: ±SEM) comparing Ctrl (E2.tdTom+saline injected intracranially) pre- and post-CNO administration (left) or hAPP (E2.tdTom+hAPP injected intracranially) pre- and post CNO administration (right). **d.** Power Spectral Density [PSD] analysis to assess if DREADD expression impacts excitability in the absence of CNO administration (dark lines: mean, light lines: ±SEM) comparing Ctrl (E2.tdTom+saline injected intracranially) to Ctrl DREADD (E2.tdTom+ E2.Gq.DREADD+saline injected intracranially) (left) or hAPP (E2.tdTom+hAPP injected intracranially) to hAPP DREADD (E2.tdTom+E2.Gq.DREADD + hAPP injected intracranially) (right).

**Extended Data Figure 15.**
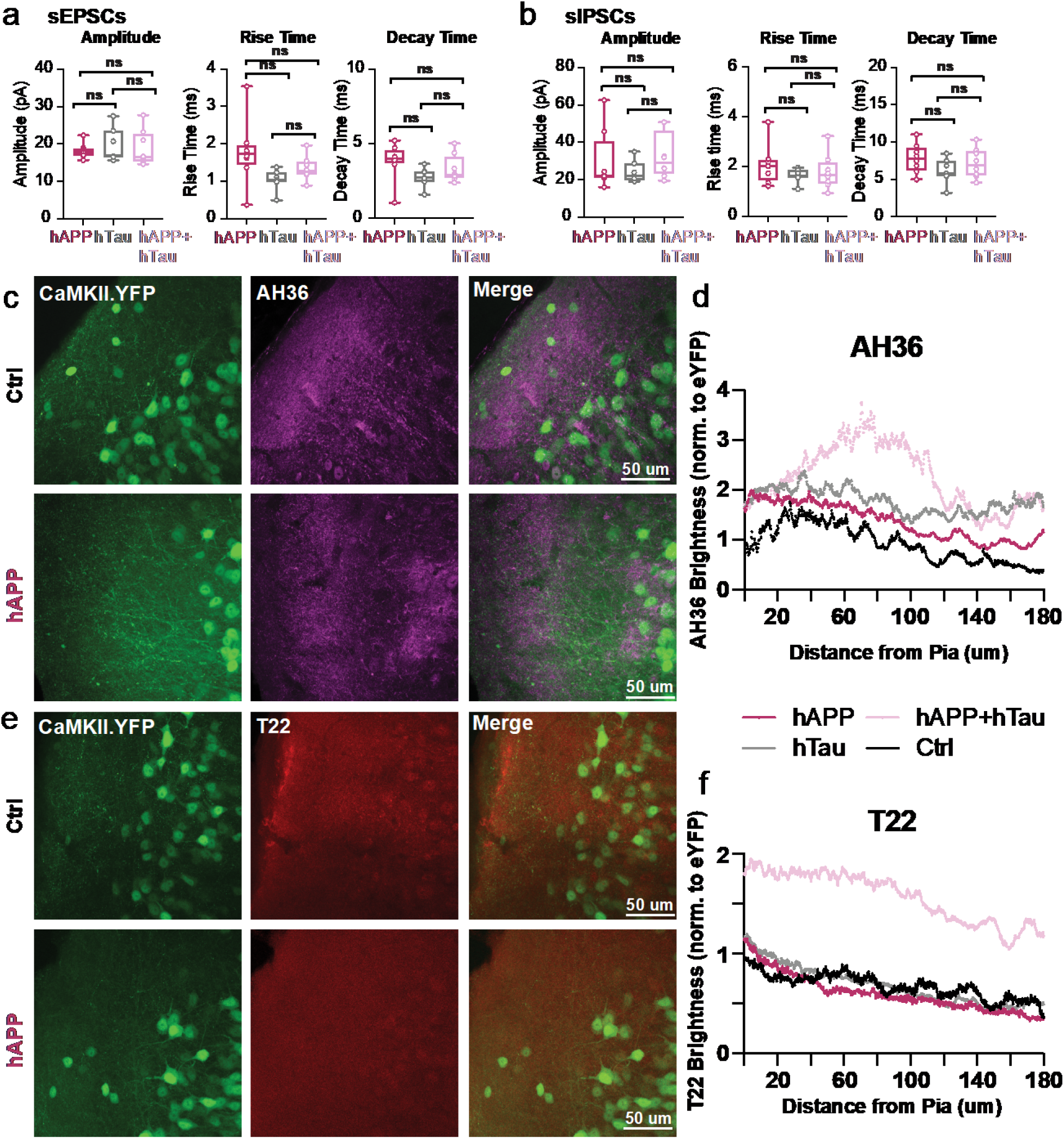
hTau co-injection with hAPP spontaneous properties and Ctrl IHC images. **a.** Summary data of sEPSC properties. sEPSC properties between hAPP injection, hTau injection, or hAPP+hTau injection are not significantly different (Amplitude: hAPP vs. hTau p=0.75, hAPP vs. hAPP+hTau p=0.94, hTau vs. hAPP +hTau p=0.92; Rise Time: hAPP vs. hTau p=0.05, hAPP vs. hAPP+hTau p=0.31, hTau vs. hAPP +hTau p=0.55; Decay Time: hAPP vs. hTau p=0.11, hAPP vs. hAPP+hTau p=0.54, hTau vs. hAPP +hTau p=0.55; df=20, One-way ANOVA with Multiple Comparisons). **b.** Summary data of sIPSC properties. sIPSC amplitudes between hAPP injection, hTau injection, or hAPP+hTau injection are not significantly different (Amplitude: hAPP vs. hTau p=0.69, hAPP vs. hAPP+hTau p=0.88, hTau vs. hAPP +hTau p=0.41; Rise Time: hAPP vs. hTau p=0.39, hAPP vs. hAPP+hTau p=0.65, hTau vs. hAPP +hTau p=0.88; Decay Time: hAPP vs. hTau p=0.21, hAPP vs. hAPP+hTau p=0.76, hTau vs. hAPP +hTau p=0.53; df=20, One-way ANOVA with Multiple Comparisons). **c,e.** IHC representative images at 60x magnification for Ctrl (top) or hAPP (bottom) injected mice with staining for either AH36 (c) or T22 (e). **d.** Ctrl, hAPP, hTau, and hAPP+hTau were analyzed for AH36 brightness using four line scans in each slice. AH36 brightness was normalized to CaMKII.eYFP brightness to control for any potential variability in viral expression. hAPP+hTau showed the highest level of AH36 brightness, most notably between 40-120 μm from the pia. **f.** hAPP, hTau, and hAPP+hTau were analyzed for T22 brightness using four line scans in each slice. AH36 brightness was normalized to CaMKII.eYFP brightness to control for any potential variability in viral expression. hAPP+hTau showed a higher level of T22 brightness, above all other groups which displayed only background levels of T22 positivity.

**Extended Data Datasheet 1. Regional-specific PV Proteomics (Attached)**

**Extended Data Datasheet 2. LEC/SS PV IntegHuman (Attached)**

**Extended Data Datasheet 3. APP and Tau Interactors (Attached)**

**Extended Data Datasheet 4. Kavanagh Tau Interactome/PV (Attached)**

